# Empathic contagious pain and consolation in laboratory rodents: species and sex comparisons

**DOI:** 10.1101/745299

**Authors:** Rui Du, Wen-Jun Luo, Kai-Wen Geng, Chun-Li Li, Yang Yu, Ting He, Na Wei, Jun Chen

## Abstract

Laboratory rodents are gregarious in nature and have a feeling of empathy when witnessing a familiar conspecific in pain. The rodent observers express two levels of empathic responses: observational contagious pain (OCP) and consolation. Here we examined the sex and species difference of OCP and consolation in male and female mice and rats. We observed no species difference in both OCP and consolation, but significant species difference in general social (allo-mouth and/or allo-tail sniffing) and non-social (self-grooming) behaviors. For sex difference, male mouse observers showed more allolicking and allogrooming behaviors toward a familiar conspecific in pain during and longer time increase in pain sensitivity after the PDSI than female mouse observers. However, no sex difference was observed in rats. Our results highlighted an evolutionary view of empathy that social animals including rodents also have the ability to feel, recognize, understand and share the other’s distressing states.

## Introduction

Increasing lines of evidence from both clinical and basic research implicate an important role of social communication in modulation of pain [1-3]. Socially coping skills among couples and family members have been suggested to relieve pain and/or negative mood under chronic conditions, probably through decreasing social stress and increasing social buffering[4-8]. Recently, some researchers indicated that pain should be redefined as a distressing experience associated with actual or potential tissue damage that involves not only sensory and emotional experience, but also cognitive and social components, highlighting the mediating roles of higher brain structures in social recognition, compassion and modulation of pain[9]. However, so far less is known about the brain processing and neural mechanisms of one’s social recognizing, understanding and sharing of suffering in pain patients due to lack of theoretical framework, animal models and experimental tools in the field of pain research and management.

Empathy for pain is a concept referred to as an evolutionary behavior of social animals and humans associated with the ability to feel, recognize, understand and share the other’s distressing (pain, social rejection and catastrophe) states through social communications and interactions[10,11]. Empathy for pain is a vicarious feeling that is felt through social transfer or contagion from a distressing object to a witnessing subject. This process has been demonstrated to be mediated by central neural network mainly consisting of the anterior cingulate cortex (ACC) and anterior insular cortex that also mediates direct emotional feeling of pain (physical pain) in humans[12-14]. Evolutionarily, witnessing distressing condition of others can motivate sympathy of a subject toward unfamiliar one, but may more deeply activate a subject’s empathic concern, consolation and desire to help toward his/her familiar social members[10,15-20], highlighting the roles of kin and group selection in development of empathy [21-23]. Meanwhile, witnessing or learning of one’s family member in pain or distress may also result in a strong feeling of pain in one’s heart through empathic contagion of pain across individuals [11,12]. Social pain associated with social rejection, defeat and failure or loss of social connections may also activate the ACC and other brain structures[24], implicating an overlap of functional neural correlates that are associated with cognition, empathy, social pain and physically emotional pain [25].

Do animals have a feeling of empathy? If yes, do animals share the same neural processing as humans do? This question is still on debate and requires to be answered by deep study and strong lines of experimental evidence. More recently, based upon the seminal discovery of reciprocal enhancement of pain across dyadic mice both in pain through social interaction[26,27], we have developed a behavioral model of empathy for pain in rats [28-30]. Experimentally, the behaviors associated with empathy for pain in rats can be at least identified as two types according to the evolutionary notion of the Russian doll model [10]. One has been referred to as an observer’s empathic consolation that is driven by social interacting with a demonstrator in pain [17,20,30,31], the other is referred to as observational contagious pain (OCP or empathic transfer of pain) from distressing object to witnessing subject[11,28-30]. Briefly, the empathic consolation in rat observers has been identified as allolicking and allogrooming behaviors toward a familiar conspecific in pain during 30 min priming dyadic social interaction (PDSI)[20,30,31]. Allolicking can be defined as an observer’s sustained licking action to a demonstrator’s injury site, while allogrooming can be defined as an observer’s head contact with the head or body of a demonstrator in pain, accompanied by a rhythmic head movement [20,30,32]. The bouts of allolicking and allogrooming behaviors can be captured by video camera recorder (VCR) and off-line analyzed qualitatively and quantitatively [32]. While, the OCP, also referred to as empathic pain hypersensitivity in our previous reports, has been identified qualitatively and quantitatively as lowered pain threshold or increased pain sensitivity in the observer rats after the PDSI with a demonstrator in pain [28-30]. The OCP remains unchanged for at least 5 h in time course measured immediately after the PDSI[29,30]. Although allogrooming behavior could be seen in both familiar and unfamiliar conspecifics during the PDSI, allolicking behavior and the OCP could only be seen in familiar observer, suggesting that the establishment of familiarity among conspecifics is essential to induction of empathic responses to other’s pain in rats [11,29,30].

Although the rat model of empathy for pain has been validated, so far the mouse model of the same paradigm has yet been reported. Moreover, whether species and sex differences exist or not for this paradigm is unknown and requires to be further addressed. Thus, to answer the above common questions, we further designed and studied the behavioral parameters associated with the OCP and consolation qualitatively and quantitatively in both male and female mice and rats.

## Methods

### Animals

Male and female C57BL/6 (B6) mice and Sprague-Dawley albino (SD) rats, purchased from the Laboratory Animal Center of the Fourth Military Medical University (FMMU), were used in this study because they represent the most frequently used laboratory rodents worldwide. Both mice and rats with age of postnatal week 4-5 were translocated from the FMMU to Tangdu Hospital SPF animal facility in which 4-6 animals of the same species and the same sex were co-housed in each cage for another 2-3 weeks so as to familiarize with each other as cagemates (Fig.1). The newly regrouped animals were fed under standard conditions with a light-dark cycle (08:00-20:00) and adjustable room temperature (25±2 °C)and air humidity (55-65%). Both water and food pellets were available *ad libitum*. This study was fully in accordance with the recommendations of the ARRIVE guidelines [33], the U.K. Animals (Scientific Procedures) Act 1986 and associated guidelines, the EU Directive 2010/63/EU for animal experiments, the National Institutes of Health guide for the care and use of laboratory animals (NIH Publications No. 8023, revised 1978), and the ethical guidelines for investigations of experimental pain in conscious animals of the International Association for the Study of Pain were also critically followed[34]. The number and suffering of animals were greatly minimized as required.

**Fig. 1.**
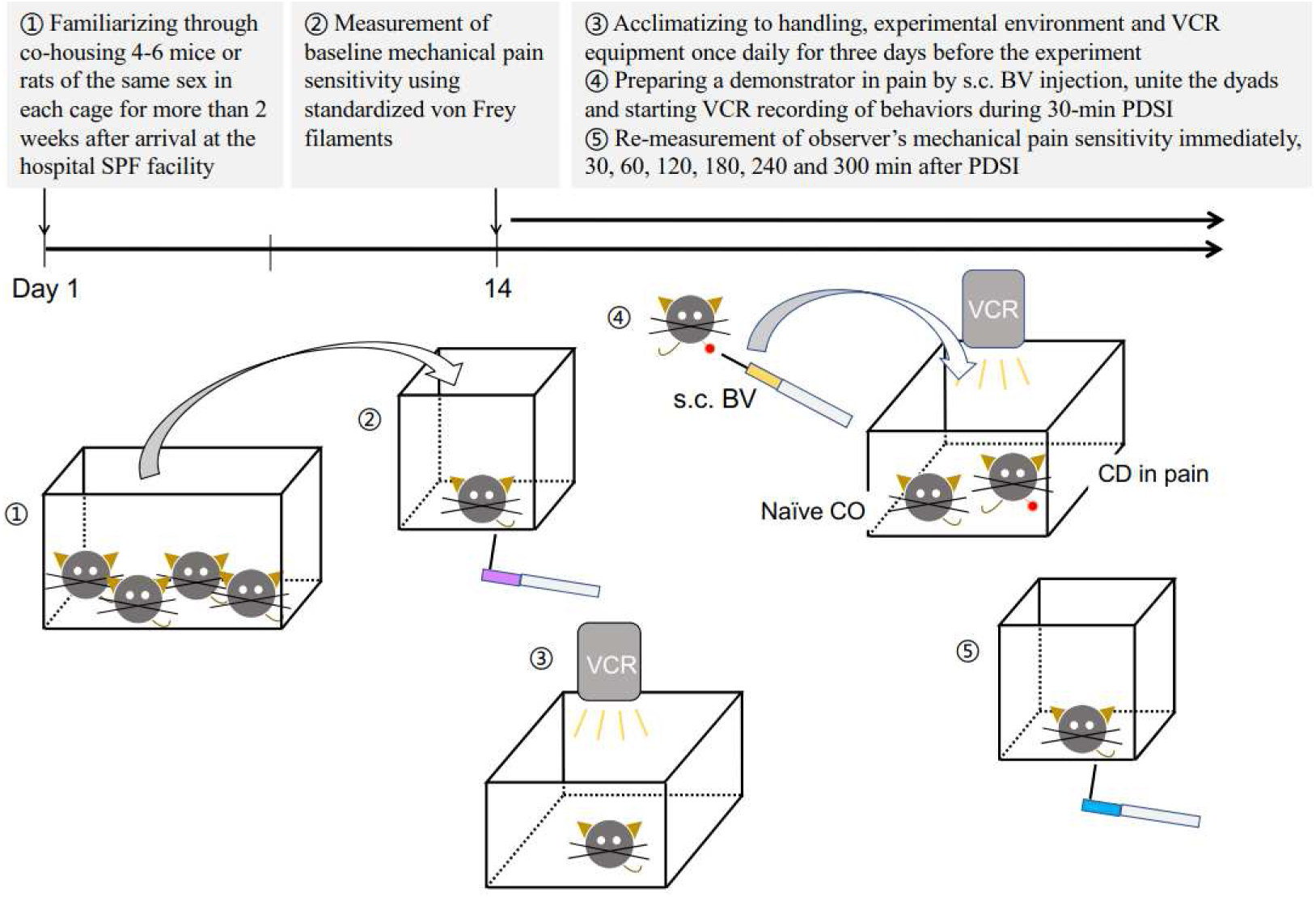
Timeline, experimental design, setup and protocol for the study of empathy for pain in mice and rats. Abbreviations: BV, bee venom; CD, cagemate demonstrator; CO, cagemate observer; PDSI, priming dyadic social interaction; s.c., subcutaneous; SPF, specific pathogen free; VCR, video camera recorder.

### Experimental design and procedures

Regardless of species, the dyads of animals used for social interaction, who were the same in sex and familiar for each other, were designed as two paradigms: (1) CO_naive_-CD_naive_, a control paradigm for dyadic social interaction between a naive cagemate observer and a naive cagemate demonstartor; (2) CO_naive_-CD_pain_, an experimental paradigm for dyadic social interaction between a naive cagemate observer and a cagemate demonstartor in pain. The timeline, design and protocol were shown in Fig.1 [32].

### Establishment of familiarity

After arrival at the hospital SPF animal facility, 4 mice or 4-6 rats of the same sex were regrouped and co-housed in each cage for more than 2 weeks (Fig.1) [32]. To avoid social conflicts among adult animals, the time for regrouping should be 3-4 weeks after birth and the number of animals to be co-housed should be limited to less than four for mice (more aggressive when stranger adults meet) and four to six for rats (less aggressive when stranger adults meet).

### Habituation to experimental procedures

The subjects to serve as an observer should be trained by acclimatizing to hand handling, experimental environment and VCR equipment once daily at least for three days before formal procedures for testing (Fig.1, for protocol details see [32]). Hand handling was a very important procedure in this study because it could buffer social stress that may block empathy for pain [27,29,30].

### Preparation of a demonstrator in pain

The selection of pain models for preparing a demonstrator in pain is another critical step for induction of empathy for pain in a witnessing observer during and after the PDSI [30,32]. As demonstrated by our previous work[30], the induction of empathy for pain in an observer rat would be determined by the observability or visibility of spontaneous pain-related behaviors displayed or expressed by a familiar demonstrator in pain. Among the animal models of pain tested, the bee venom (BV) test, the formalin test and the acetic acid test that can induce long-term robust spontaneous pain-related behaviors such as paw flinching, paw licking and lifting or abdominal writhing have been demonstrated to be effective to induce both consolation and the OCP, whereas, the complete Freund’s adjuvant (CFA) and the spared nerve injury (SNI) models that induce less spontaneous pain-related behaviors are not effective in this paradigm [30,31]. Since the BV test is both a scientifically well-established and human-rodent co-experienced type of pain[35-38], it was used in the whole experiment of this study. Briefly, the demonstrator received a subcutaneous (s.c.) injection of BV solution (25 μl for mice and 50 μl for rats, 0.4% lyophilized whole venom of *Apis millifera* dissolved in physiological saline) into the left hind paw just before the start of the VCR recording of the PDSI and then re-united with the naive observer in the testing box (for details see [32]).

### Quantitative sensory test with von Frey filaments

Because our previous studies have only identified contagious mechanical pain hypersensitivity, but not thermal hypersensitivity following the PDSI[28-30], only mechanical pain sensitivity was examined in the current study. The mechanical pain sensitivity test setting includes a supporting platform and a nontransparent plastic testing box (10.5 cm x 10.5 cm x 15.8 cm) that is necessary to prevent any visual information from coming during testing. The supporting platform (160 x 30 x 40 cm) is equipped with metal mesh. The pore size of the mesh (0.5 cm x 0.5 cm) is preferably such that both mice and rats can move freely on the surface without getting caught. Because the mechanical pain sensitivity for paw withdrawal reflex was quite different between mice and rats, different quantitative method was used in this study. For both mice and rats, the mechanical pain sensitivity of the observer was measured prior to (1 day before for baseline) and after the PDSI (immediate, 30, 60, 120, 180, 240, 300 min). For mice who are likely to have high mechanical pain sensitivity and more active in motion in nature, an ascending series of calibrated von Frey (vF) filaments with intensities ranging from 0.16 to 1.40 g (1.60 to 13.72 mN) were used to induce paw withdrawal reflex from minimum (0) to maximum (100%). With the increasing intensity, each stimulus should be continued 1-2 s for 5 repetitions in 5 s apart, avoiding the same site. A sharp withdrawal or lift up after a stimulus was considered a positive response and should be recorded. The averaged percent response (%) of a mouse to 5 stimuli of each intensity was calculated and pooled into stimulus-response functional curves (SRFC). Comparing to the baseline, leftward shift of the SRFC was defined as hypersensitivity (hyperalgesia or allodynia), while rightward shift of the curve was defined as hyposensitivity (analgesia)[39]. Finally, the fitted vF intensity of half maximal response was obtained by Bliss method[40], serving as relative mechanical threshold for mice. For rats who have relatively low mechanical pain sensitivity and inactive in motion in nature, a series of calibrated vF filaments with bending force intensities ranging from 2.00 to 60.00 g (19.60 to 588.00 mN) were used to induce paw withdrawal reflex. The bending force of a vF filament that enabled 50-60% response to 10 stimuli was determined as the paw withdrawal mechanical threshold (PWMT). For details see our published protocol [32].

### PDSI and VCR recording

The PDSI has been defined as a preemptive condition that allows full body contact, social communication and interaction between a naive observer and a demonstrator for 30 min [11]. A naive observer meant that the subject animal had no experience of pathologically tissue or nerve injury at all but only had experienced physiologically stroking stimulus by vF filaments one day before the PDSI[11]. Briefly, a VCR (Sony, FDR-AX40, Japan) setting was arranged in a right top-down vertical view over the testing box(19 x 19 x 30 cm for mice and 40 x 30 x 15 cm for rats) which was used as an arena for 30 min PDSI (for details see [32]).

### Offline qualitative identification and quantitative analyses of social and non-social behaviors during PDSI

According to repeated observations of the VCR-based behaviors in a 30 min lapse of time, the behaviors were classified into three types: (1) consolation behavior identified as allolicking and allogrooming that has been believed to be reciprocal altruistic behaviors [16,17,20,30]; (2) general social behaviors identified as allo-mouth and/or allo-tail sniffing[41]; (3) non-social behavior identified as self-licking and self-grooming that is an innate stereotyped and patterned behavior of rodents and other terrestrial mammals generated and controlled by the brain [16,17]. For each type of behaviors, the latency for the observer subject to first perform a type of behaviors after initiation of the PDSI, the time course and total time the observer subject spent on a type of behaviors during 30 min period of PDSI, and the total counts the observer subject visited and behaved for each type of behaviors during 30 min period of PDSI were rated and statistically analyzed. Both social and non-social behaviors were captured by the VCR in real time, and qualitatively identified and quantitatively analyzed offline by one to two analyzers who were blind to the treatment of animals. Grooming of less than 1 s was excluded. Grooming directed toward the genitals was excluded in this study.

### Statistical analysis

All data were presented as mean ± SEM. SPSS 25.0 was used for data analyses that could perform automatically overall corrections for various statistical tests used. In principle, parametric statistical analysis methods would be used if both normality test and equal variance test for samples were passed, however, only non-parametric statistical analysis method would be used if either of the normality test or equal variance test failed (for details see Table SI). Normality of the distribution was analyzed by Shapiro-Wilk test, while homogeneity of variance was analyzed by Levene test. Nonparametric two-tailed Mann-Whitney *U* test or parametric two-tailed *t*-test were used depending upon the results of the normality and homogeneity tests. Two-way ANOVA repeated measure (RM) with Bonferroni post hoc correction was used for time course data. For within-time two-way ANOVA RM, Greenhouse-Geisser method was used if Mauchly’s test of sphericity failed. For paired comparisons, Wilcoxon Signed Rank Test, Friedman′s *M* test and Mann-Whitney *U* test (two-tailed) were used if Shapiro-Wilk test and Equal variance test failed (for details see Tables SII-SIII). *P*< 0.05 was considered as statistically significant. Graphs and plots in the illustrations were made by GraphPad Prism version 7.0a.

## Results

### Species and sex comparisons of consolation behavior

The observers from the CO_naive_-CD_pain_ paradigm of both mice and rats showed more consolation behaviors toward the conspecific in pain when comparing with the observers from the CO_naive_-CD_naive_ paradigm (TablesI-II). Generally, both mouse and rat observers had shorter latency and more time and visit counts engaged in allolicking and allogrooming when witnessing a conspecific in pain (TablesI-II). Interestingly, the mouse observers also had allo-mouth sniffing behavior, however, in contrast the rat observers did not have any allo-mouth sniffing behavior (TablesI-II and Figs.2-3). Both mouse and rat observers had allo-tail sniffing and self-licking/self-grooming behaviors as previously described (TablesI-II and Figs. 2-3).

**Table I.**
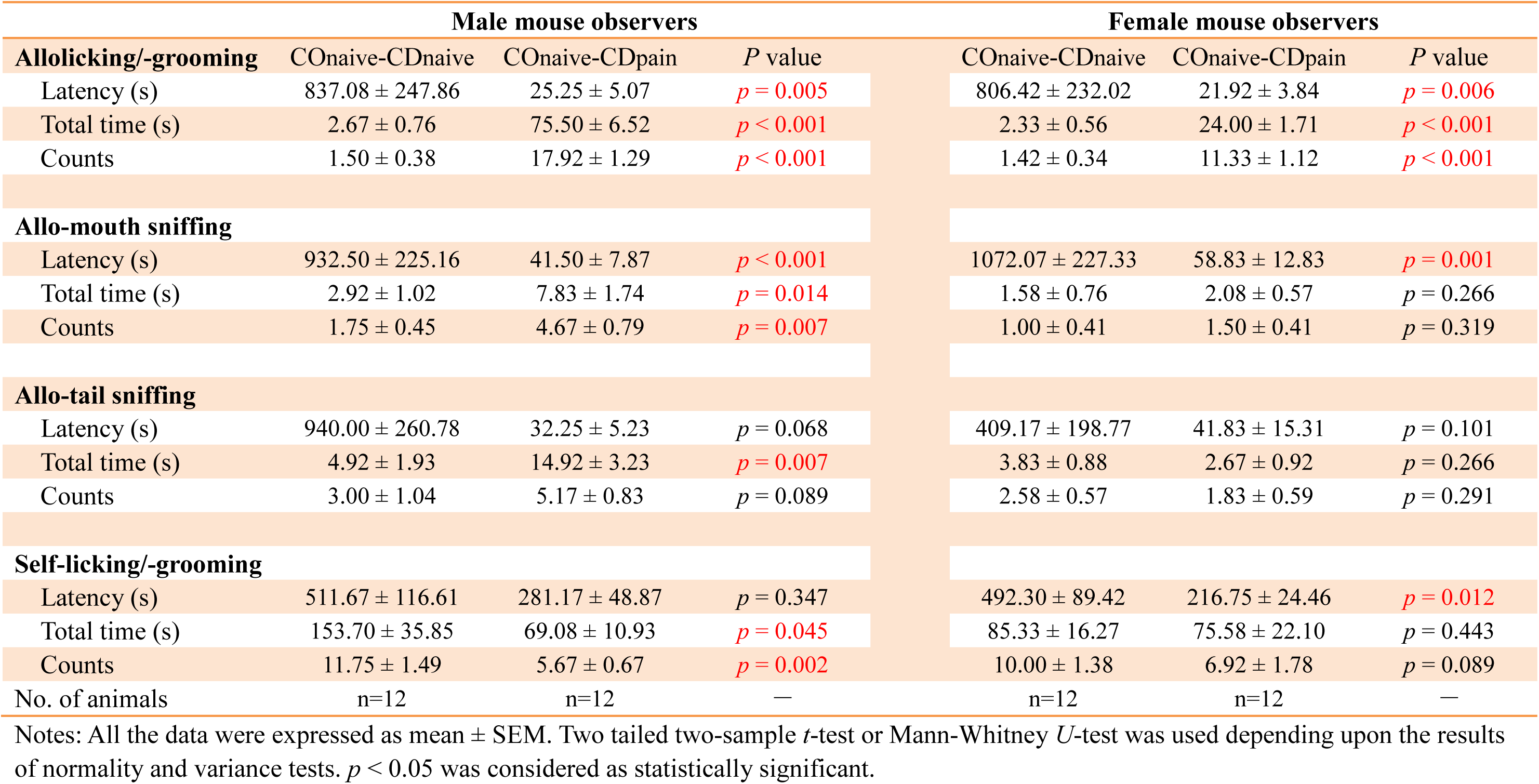
Comparisons of empathic consolation, general social and non-social behaviors among the cagemate observers of COnaive-CDnaive and COnaive-CDpain in mice of the same sex

**Table II.**
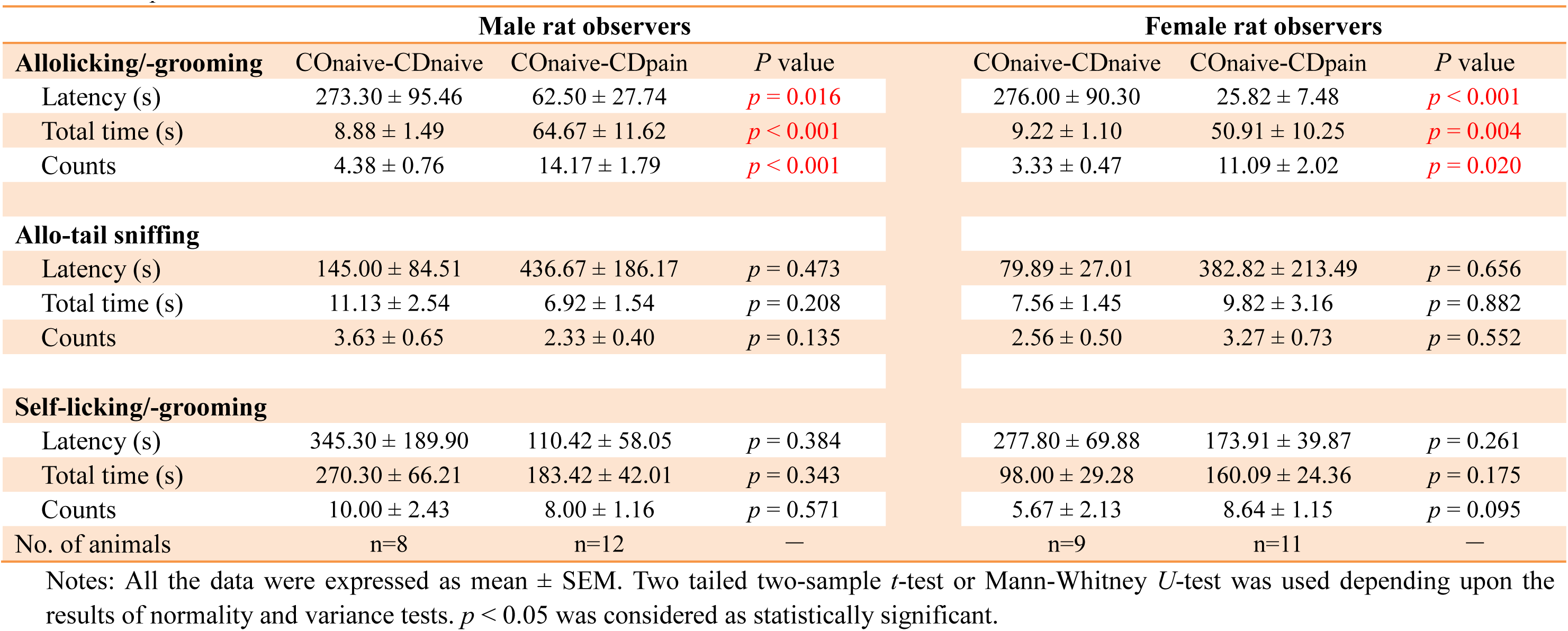
Comparisons of empathic consolation, general social and non-social behaviors among the cagemate observers of COnaive-CDnaive and COnaive-CDpain in rats of the same sex

**Fig. 2.**
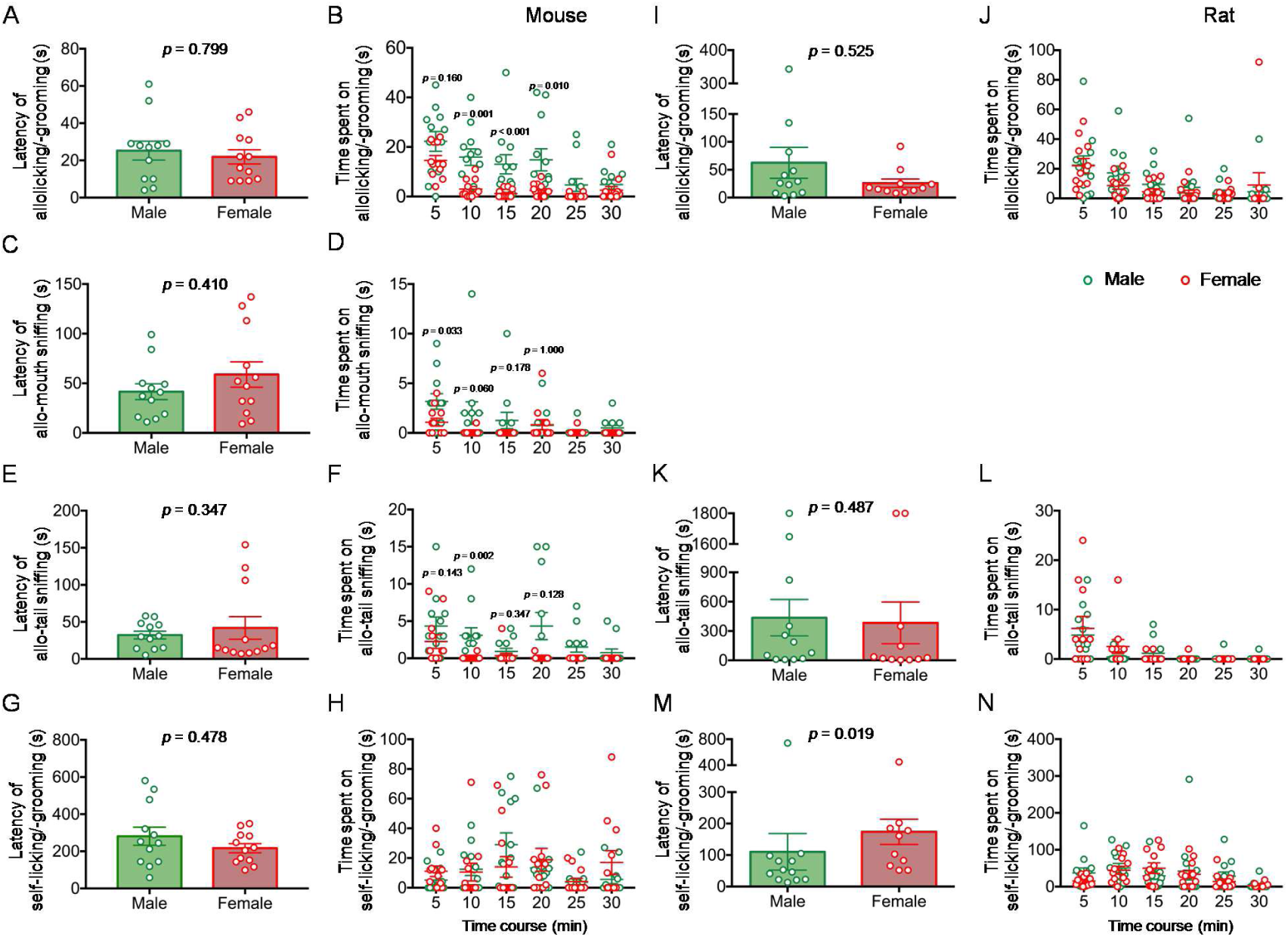
Sex and species comparisons of consolation (allolicking/allogrooming), general social (allo-mouth and/or allo-tail sniffing) and non-social (self-licking/self-grooming) behaviors between male and female observers in mice (A-H) and rats (I-N) during 30-min priming dyadic social interaction with a cagemate demonstrator of the same sex in pain. Latencies and time courses engaged by the cagemate observer in allolicking/allogrooming (**A-B** for mice and **I-J** for rats), allo-mouth sniffing (**C-D** for mice), allo-tail sniffing (**E-F** for mice and **K-L** for rats) and self-licking/self-grooming (**G-H** for mice and **M-N** for rats). *p* < 0.05 as statistical significance [Male (n=12) *vs*. Female (n=11-12) for each species, with two-tailed two-sample *t*-test or Mann-Whitney *U* test, for details see **Table SI-SII**]. Mean±SEM.

**Fig. 3.**
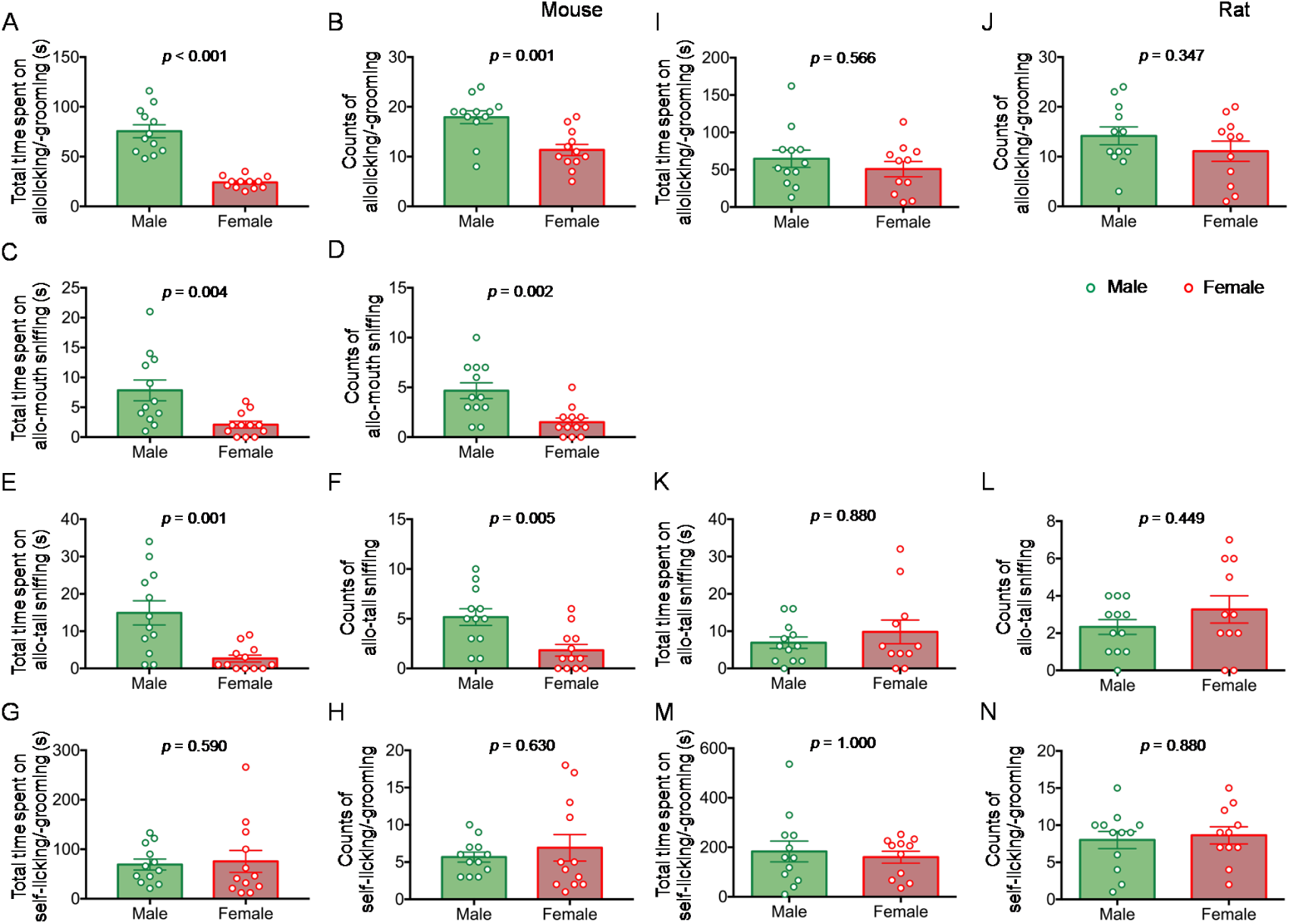
Sex and species comparisons of consolation (allo-licking/allo-grooming), general social (allo-mouth and/or allo-tail sniffing) and non-social (self-licking/self-grooming) behaviors between male and female observers in mice (A-H) and rats (I-N) during 30-min priming dyadic social interaction with a cagemate demonstrator of the same sex in pain. Total time and visit counts spent by the cagemate observer on allolicking/allogrooming (**A-B** for mice and **I-J** for rats), allo-mouth sniffing (**C-D** for mice), allo-tail sniffing (**E-F** for mice and **K-L** for rats), and self-licking/self-grooming (**G-H** for mice and **M-N** for rats). *p*< 0.05 as statistical significance [Male (n=12) *vs*. Female (n=11-12) with two-tailed Mann-Whitney *U* test, for details see **Table SI**]. Mean±SEM.

#### Species comparisons of consolation behavior

There was no species difference in latency, total time and counts of allolicking and allogrooming between B6 mice and SD rats in either male or female (Table III). Species difference was not revealed in allo-tail sniffing in terms of latency and total time between mice and rats in either male or female (Table III). Although male mice had more counts than male rats (*p* = 0.002, Mann-Whitney *U* test), species difference was not seen between mice and rats of female for the counts of allo-tail sniffing (Table III). As for the non-social behavior, rats of both sexes spent more time in self-licking and self-grooming than mice of both sexes (Table III, mice vs. rats: *p* = 0.017 for male and *p* = 0.016 for female, Mann-Whitney *U* test) although counts showed no species difference. Moreover, rats of both sexes had shorter latency in self-grooming than mice of both sexes although statistical significance for species difference was only seen in male (Table III, *p* = 0.001, Mann-Whitney *U* test).

**Table III.** Cross-species comparisons of empathic consolation, general social and non-social behaviors in subjects of the same sex.

|  | Male observer |  |  | Female observer |  |  |
| --- | --- | --- | --- | --- | --- | --- |
|  | Mouse | Rat | *P* value | Mouse | Rat | *P* value |
| **Allolicking/-grooming** |  |  |  |  |  |  |
| Latency (s) | 25.25 ± 5.07 | 62.50 ± 27.74 | *p* = 0.630 | 21.92 ± 3.84 | 25.82 ± 7.48 | *p* = 0.880 |
| Total time (s) | 75.50 ± 6.52 | 64.67 ± 11.62 | *p* = 0.425 | 24.00 ± 1.71 | 50.91 ± 10.25 | *p* = 0.079 |
| Counts | 17.92 ± 1.29 | 14.17 ± 1.79 | *p* = 0.143 | 11.33 ± 1.12 | 11.09 ± 2.02 | *p* = 0.928 |
| **Allo-tail sniffing** |  |  |  |  |  |  |
| Latency (s) | 32.25 ± 5.23 | 436.67 ± 186.17 | *p* = 0.101 | 41.83 ± 15.31 | 382.82 ± 213.49 | *p* = 0.347 |
| Total time (s) | 14.92 ± 3.23 | 6.92 ± 1.54 | *p* = 0.089 | 2.67 ± 0.92 | 9.82 ± 3.16 | *p* = 0.051 |
| Counts | 5.17 ± 0.83 | 2.33 ± 0.40 | *p* = 0.006 | 1.83 ± 0.59 | 3.27 ± 0.73 | *p* = 0.151 |
| **Self-licking/-grooming** |  |  |  |  |  |  |
| Latency (s) | 281.17 ± 48.87 | 110.42 ± 58.05 | *p* = 0.001 | 216.75 ± 24.46 | 173.91 ± 39.87 | *p* = 0.190 |
| Total time (s) | 69.08 ± 10.93 | 183.42 ± 42.01 | *p* = 0.017 | 75.58 ± 22.10 | 160.09 ± 24.36 | *p* = 0.016 |
| Counts | 5.67 ± 0.67 | 8.00 ± 1.16 | *p* = 0.095 | 6.92 ± 1.78 | 8.64 ± 1.15 | *p* = 0.260 |
| No. of animals | n=12 | n=12 | — | n=12 | n=11 | — |

Notes: All the data were expressed as mean ± SEM. Two tailed two-sample *t*-test or Mann-Whitney *U*-test was used depending upon the results of normality and variance tests. *P* < 0.05 was considered as statistically significant.

Taking the data of latency and time course together (Figs. 2 and 3), it was revealed that both mouse and rat observers of either male or female were likely to approach to the conspecific in pain as quickly as possible and spent more time on consolation and social behaviors than self-grooming behavior.

#### Sex comparisons of consolation behavior

In mice, sex difference was distinctly seen in both empathic consolation and general social behaviors in terms of time and counts but with latency being of no sex difference (Fig.2A-H and Fig. 3A-H, see supplemental Tables SI-SII for statistical analysis). Male mice spent more time and had more visit counts than female in allolicking/allogrooming and allo-mouth/allo-tail sniffing toward a conspecific in pain during the early 20 min PDSI (Fig.2B, Fig. 2D, Fig. 2F, Fig. 2H, see supplemental Tables SI-SII for statistical analysis), while there was no sex difference in self-licking/self-grooming in terms of latency, time and counts (Fig.2A-H and Fig. 3A-H, see supplemental Tables SI-SII for statistical analysis).

In rats, no sex difference was seen in either empathic consolation or general social behavior in terms of latency, time and counts (Fig.2I-L and Fig. 3I-L, see supplemental Tables SI-SII for statistical analysis). Although female rats likely had relatively shorter latency than male (*p* = 0.019, Mann-Whitney *U* test), no sex difference was seen in time and counts of self-grooming behavior (Fig.2M-N and Fig. 3M-N, see supplemental Tables SI-SII for statistical analysis).

### Species and sex comparisons of observational contagious pain

Similar to our previous reports on rats (also see Fig.5C-F)[28-30,32], the OCP occurred as well in naive mouse observers from the CO_naive_-CD_pain_ paradigm after 30 min PDSI (Fig. 4G-L and Fig. 5B), whereas, the OCP could not be identified in the mouse observers from the CO_naive_-CD_naive_ paradigm (Fig.4A-F and Fig. 5A). Both male and female mouse observers presented long-term mechanical pain hypersensitivity after the PDSI with a conspecific in pain, being evidenced by significant leftward shift of the SRFC from the baseline (Fig. 4G-L, see supplemental Tables SI-SIII for statistical analysis) and lowered PWMT (fitted vF intensity for half maximal response, see Fig. 5B). The OCP identified in the mouse observers from the CO_naive_-CD_pain_ paradigm did not disappear until 240 min in female and 300 min in male after the PDSI (Fig.4G-L and Fig. 5B, see supplemental Tables SI-SIII for statistical analysis).

**Fig. 4.**
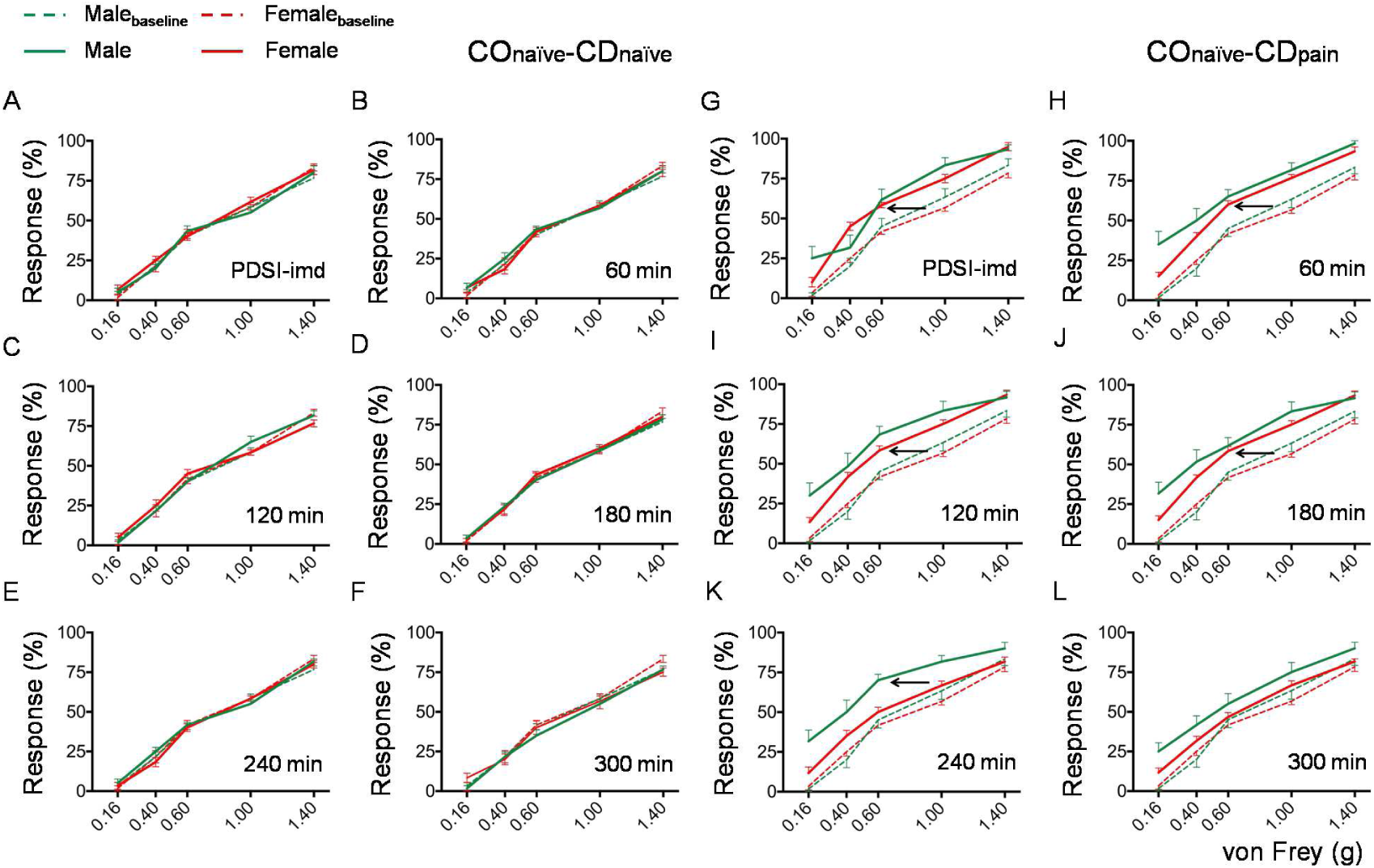
Sex comparisons of the stimulus-response functional curves for mechanical pain sensitivity in mouse observers from the CO_naive_-CD_naive_ and the CO_naive_-CD_pain_ paradigms prior to (Baseline, dashed) and immediately (PDSI-imd, **A, G**), 60min (**B, H**), 120min (**C, I**), 180min (**D, J**), 240min (**E, K**) and 300min (**F, L**) after priming dyadic social interaction (PDSI) with a cagemate demonstrator of the same sex in pain. *p*< 0.05 as statistical significance [Male (n=12) *vs*. Female (n=12) with two-tailed Mann-Whitney *U* test, for details see **Table SII-SIII**]. BL, baseline. Mean±SEM.

**Fig. 5.**
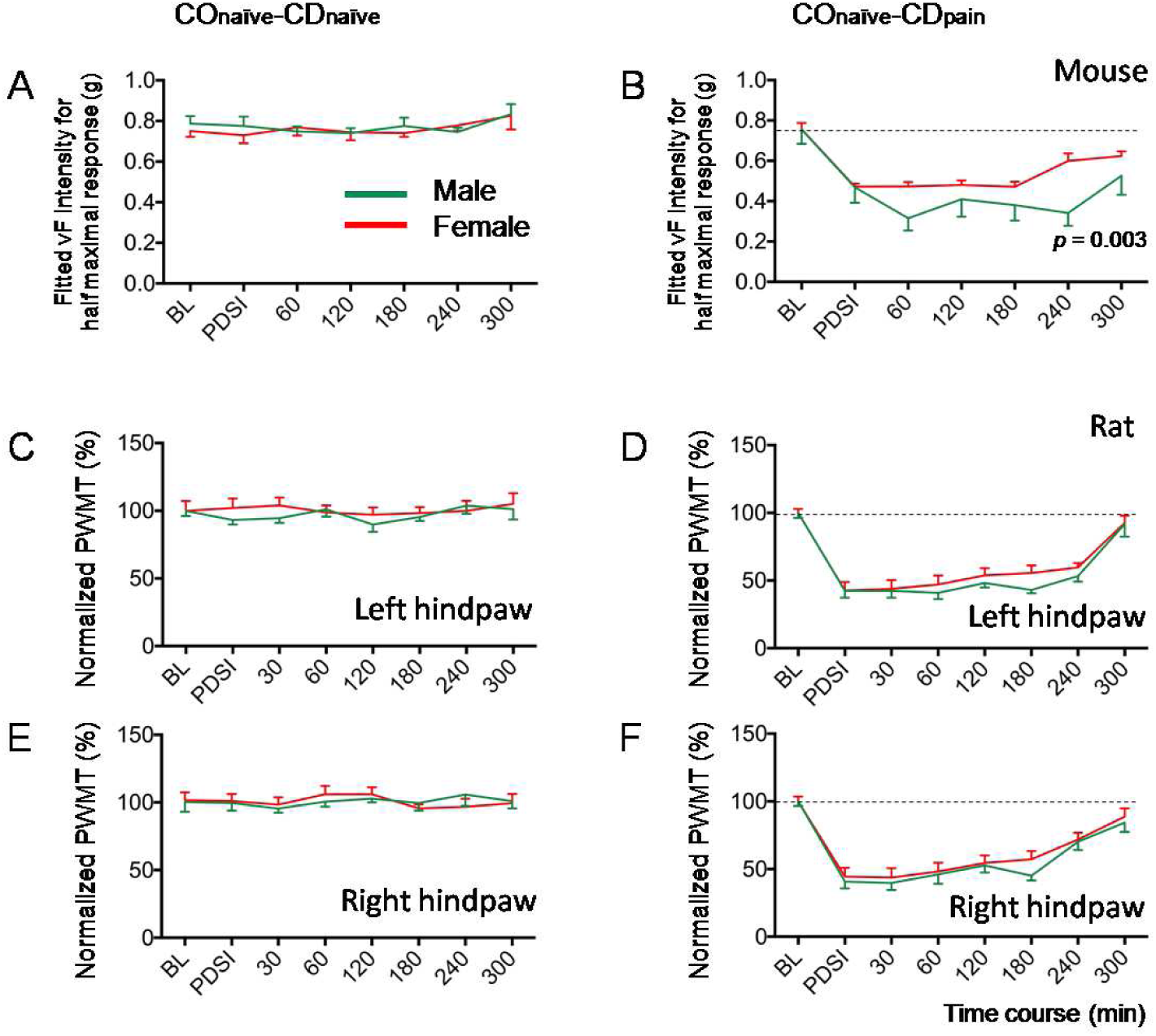
Sex comparisons of changes in mechanical pain sensitivity in mouse and rat observers from the CO_naive_-CD_naive_ and the CO_naive_-CD_pain_ paradigms prior to (BL) and immediately, 30, 60, 120, 180, 240 and 300 min after priming dyadic social interaction (PDSI) with a cagemate demonstrator of the same sex in pain. (A-B) Time courses of changes in von Frey (vF) intensity (g) for half maximal response fitted from the stimulus-response functional curves of **Figure 4** in mouse observers by Bliss method. (C-F) Time courses of normalized paw withdrawal mechanical threshold (PWMT) measured in the left (C-D) and right (E-F) hindpaws of rat observers. BL, baseline. *p*< 0.05 as statistical significance [Male (n=8-12) *vs*. Female (n=12) with two-tailed Mann-Whitney *U* test, for details see **Table SII-SIII**]. Mean±SEM.

#### Species comparisons of observational contagious pain

Generally speaking, no species difference in the OCP was revealed between mice and rats of either sex in terms of magnitude and time course under the same experimental condition, procedure and paradigm(Fig. 5B, Fig. 5D, Fig. 5F, see supplemental Tables SI-SIII for statistical analysis).

#### Sex comparisons of observational contagious pain

No sex difference was revealed in the OCP between male and female observers in either mice or rats in terms of magnitude and time course under the same experimental condition, procedure and paradigm between 0-180 min period after the PDSI (Fig. 5B, Fig. 5D, Fig. 5F, see supplemental Tables SI-SIII for statistical analysis). However, the empathic mechanical pain hypersensitivity in mouse observer was maintained relatively longer in male than female (Fig. 4G-L and Fig. 5B, see supplemental Tables SI-SIII for statistical analysis). No sex difference was seen in the OCP between male and female observers in rats during the whole time of observation (Fig. 5D and Fig. 5F, see supplemental Tables SI-SII for statistical analysis).

## Discussion

### Evidence for evolutionary issue of empathy

From the point of evolutionary view, empathy has been proposed to be hierarchical in mammals that has evolved from very low stage (motor mimicry and emotional contagion) to relatively higher stage (empathic concern and consolation), and finally to the highest stage (perspective-taking, mentalizing, theory of mind and targeted-help) from lower animals to human beings [10]. Although several emerging lines of evidence support existence of emotional contagion in lower mammals[3,11,41-43], answers to the questions about whether lower mammals are able to recognize, understand, share and care others are still controversial due to lack of enough direct experimentally supporting evidence [20,30,44]. In a series of reports associated with empathy for pain in rats and mice including the present study, our lab has provided with strong lines of experimental evidence supporting existence of both emotional contagion and empathic consolation in laboratory rodents[11,28-32]. Before the coming of our findings, empathic consolation has only been observed in a special sub-species of wild rodents - socially monogamous, biparental prairie vole [20] although emotional contagion of pain or observational fear learning have been increasingly evidenced [3,10,11]. Taken together, it has been demonstrated experimentally that lower mammals such as rodents may have both lower stage (emotional contagion, i.e., OCP here) and relatively higher stage (empathic concern and consolation) of empathy, supporting the rationality of theoretical Russian-doll model for the evolution of empathy in mammals [10]. Moreover, the findings that social familiarity plays essential roles in induction of empathy for pain in rodents also support Darwin’s assertion that “with all animals, sympathy is directed solely towards the members of the same community, and therefore towards known, and more or less beloved members, but not to all the individuals of the same species”[11,15] and the theories of kin or group selection [21-23].

### Qualitative and quantitative assessment of empathy for pain in laboratory rodents

In the past century, study of empathy has been mostly performed in non-human primates and other non-laboratory animals outdoors [10,41-43]. This has greatly limited the number of researchers joining the study and hindered the advances of empathy research in terms of bio-psychosocial-brain-behavioral paradigm [11,41-43]. Therefore, discovering, developing and validating the laboratory animal models of empathy would be very important and critical for opening a new field of science - neuroscience of empathy. Here we have developed a laboratory rodent model of empathy for pain in both mice and rats using a set of novel behavioral parameters for both qualitative and quantitative assessment. We have identified and validated two behavioral identities of empathy for painfrom laboratory rodent model: (1) consolation; and (2) observational contagious pain.

#### Are there species and sex differences in consolation between mice and rats?

To make qualitative and quantitative assessment of consolation, we successfully identified allolicking and allogrooming behaviors from the naive observers during PDSI with a familiar conspecific in pain. To see whether the observer’s allolicking and allogrooming behaviors are selective or specific to the injury and pain of the object, we also evaluated general social behavior (allo-mouth and allo-tail sniffing) and non-social behavior (self-licking and self-grooming) in the observers. In each type of targeted behaviors, four bio-parameters including latency, time course, total time and visit counts were quantitatively assessed. In the present study, it was clearly shown that there was no species difference between mice and rats for allolicking and allogrooming behaviors in either male or female, suggesting laboratory rodents can be motivated to perform empathic consolation when witnessing their familiars in painful or distressing condition. Mice and rats are likely sharing and caring as humans. Bio-parameter data showed that both mouse and rat observers began to approach toward the familiar conspecific in pain in a short delay when witnessing and then spent longer time to allolick the injury site and to allogroom the body of the injured partners. As contrast, the same animals had longer latency and less visit count in either self-licking/self-grooming or allo-tail and allo-mouth sniffing, suggesting that laboratory rodents have a strong ability to rapidly recognize and understand the distressing condition of others. And this process is likely to motivate visiting, sharing and caring of the injured object at the expense of loss of their time in environmental exploration and self-grooming. Because self-grooming is predominant in rodents’ usual behaviors (more than 40% of living time) [16,17], loss of self-grooming and gain of allolicking and allogrooming in time during PDSI highly implicate existence of prosocial and altruistic behaviors in observer rodents while witnessing a familiar in pain.

It is interesting to note that there was a sex difference in visit counts and total time of allolicking and allogrooming as well as allo-mouth and allo-tail sniffing between male and female mice, however, no such sex difference was seen in rats. Unlike the results from humans and rodents that female are more empathic than male [45-47], in the current study, however, the male was likely to spend more time (three folds) than the female in mice to allolick and allogroomthe injured partner. Although the female mouse observers had less time engaged in allogrooming but spent more time on allolicking toward the BV-induced injury site in the distressing object, the sex difference in empathic consolation in mice is not likely to be only caused by the sex difference in allogrooming since general social behaviors (allo-mouth and allo-tail sniffing) also had sex difference. Generally, the male has more consolation and more social behaviors than the female in mice. Moreover, rats had equivalent amount of time and visit chance in allolicking, allogrooming and allo-tail sniffing between male and female. Unlike mice, the rat observers showed less time engaged in allo-mouth sniffing although allo-tail sniffing had no difference in time between the two species. Although the underlying mechanisms of sex difference in the degree of empathic consolation and general social behaviors in mice are not clear, the level of sex hormones, genetic background and other unknown factors should be considered. In mice, for instance, variability in empathic fear response has already been noted across different inbred strains [48,49]. Because we only compared two species of laboratory rodents in which B6 mouse is one of many inbred strains and the SD rat is outbred, comparisons among different strains of inbred mice might be more significant for identification of genetic basis of empathy for pain.

#### Are there species and sex differences in observational contagious pain between mice and rats?

As aforementioned, although mice and rats have different mechanical sensitivity to vF stimuli, standardized measurements revealed no species and sex differences in observational contagious pain. Similar to our previous reports on male rats[29,30], the current data further showed that the rat observers had no sex difference in empathic mechanical pain hypersensitivity between male and female after PDSI with a familiar conspecific in pain. The paw withdrawal threshold of both sexes became lowered by more than 50% immediately after the PDSI, and the lowered mechanical threshold was maintained unchanged until 300 min of the observation. The relative long-term decrease in mechanical threshold could be identified in both sides of the hind paws and was in parallel between male and female in the rat observers. Similarly, empathic mechanical pain hypersensitivity was also identified in the mouse observers of both sexes immediately after the PDSI by showing leftward shift of the SRFC from the baseline curves. The leftward shift of the SRFC remained unchanged between male and female mice until 240 min after the PDSI. Moreover, the fitted vF intensity for the half maximal response in mice that is equivalent to the PWMT in rats also showed a separation of time effect between male and female at 240 min after the PDSI. Because the male observers had longer time course in both the consolation and the OCP than the female did, this may reflect a higher correlation between the two empathic behaviors in mice. Although sex- and gender-difference in pain have been well established in human beings [50], the sex-difference in empathic contagious pain in mice is not likely to be attributed to the sex-difference in mechanical pain sensitivity because the SRFC for the observers from the CO_naive_-CD_naive_ paradigm (for both pre- and post-PDSI) and the baseline from the CO_naive_-CD_pain_ paradigm overlapped well between male and female.

### Laboratory rodent model of empathy for pain and its advantages in application

Based upon the present results from species and sex studies, male rats and mice are highly recommended to be used as observer subjects for study of empathy for pain in laboratory rodents due to alterations of empathic responses in female mice. Because female are more sensitive to pain stimuli and more susceptible to chronic pain conditions than male in both human and animal subjects due to biopsychosocial variables [50], pain mechanisms in female are also more complex than male. Moreover, familiar conspecifics of the same sex for PDSI are also recommended because sexual behaviors could not be completely excluded if heterosexual cagemates were allowed.

As introduced in our previous reports, the selection of pain models for preparing a demonstrator in pain is also important and critical. The more visually distinctly visible the pain-related behaviors displayed by the object in pain, the more empathic responses could be induced in the rat observers in terms of both the OCP and consolation [30]. Namely, rat observers showed more consolation (allolicking and allogrooming) behaviors during PDSI with a conspecific treated with BV than with CFA[30]. Meanwhile, rat observers have distinct empathic contagious pain after PDSI with a conspecific treated with BV and formalin but do not have empathic contagious pain after the same period of PDSI with a conspecific prepared with CFA and spared nerve injury [30]. These results suggest important roles of visual information in the induction and maintenance of the OCP and consolation as suggested by a previous report [26].

The advantages of the laboratory rodent (rats and mice) model of empathy for pain are as follows: (1) unlike wild animals such as prairie voles, laboratory rodents are fed in a SPF animal facility and tested in a standardized experimental environment that are safe in prevention of infectious disease transmission from animal to animal and from animal to experimenters; (2) biological control makes genetic background of laboratory rodents more clear and comparable than wild animals; (3) attracting and recruiting more biologists and neuroscience researchers who are interested in biological basis of empathy to join us in the empathy research; (4) unlike the “double pain paradigm” introduced by Mogil’s lab [26], the laboratory rodent observers used in our paradigm are under naive condition prior to and during PDSI that can completely exclude the distressing effects of tonic pain stimulation on the observers themselves and make neurobiological, endocrine and other biological assays possible in further tests; (5) the laboratory rodent model of empathy for pain has been validated to have both empathic consolation and empathic contagious pain that are useful paradigms for studying evolutionary issues of empathy in mammals [10,11]; (6) our laboratory rodent model of empathy for pain has been approved to be mediated by top-down facilitation from the medial prefrontal cortex and the locus coeruleus-norepinephrine system [28,29] that are known to be also important brain structures involved in empathy for pain in humans [12,51]; (7)our laboratory rodent model of empathy for pain will provide a novel bio-psychosocial-brain-behavioral paradigm that can be used in combination with other advanced techniques in neuroscience such as optogenetic, chemogenetic, *in-vivo* multi-electrode array recordings and other neuroimaging approaches in consciously socially interacting animals.

## Supporting information

Supplemental Table SII

Supplemental Table SIII

Supplemental Table SIV

## Acknowledgments

The authors are grateful to Y.-Q. Yu, W. Sun, Y. Wang, Y.-J. Yin, R.-R. Wang, Y. Yang and F. Yang for cooperation and X.-L. Wang for animal support. This work was supported by grants from the National Natural Science Foundation of China to J. C (81571072) and to C.-L. L (31600855).

## Author contributions

R. Du and W.-J. Luo designed and performed the mouse and rat experiments, respectively, participated in data analysis and graph plotting. K.W. Geng contributed to the setup of VCR and initial identification of empathic consolation in mice. T. He performed the partial male mouse experiments. C.-L. Li, Y. Yu, N. Wei, and T. He provided recommendations for experimental design and participated in some behavioral quantitative analyses. J. Chen managed the whole procedure, integrated the whole data (illustrations and tables) and wrote the manuscript. All authors participated in the discussion of the results and the final revision of the manuscript.

## Supplemental materials

Table SI. Detailed descriptions of the number of animals used and statistical analyses for each part of the experiments.

Table SII. Time effects of empathic consoling and empathic contagion of pain in mice and rats of both sexes.

Table SIII. Sex comparisons of stimulus-response functional curves in mice. Table SIV. Sample size prediction by one-way ANOVA power analysis.

