## Supplemental Table SI for "Empathic contagious pain and consolation in laboratory rodents: species and sex comparisons"

Table SI Detailed descriptions of the number of animals used and statistical analyses including normality test and equal variance test for samples, statistical methods and *p* or *U/t* values for each part of the experiments shown in Figs 2-5.

| Figure number | No of animals | Normality test | Equal variance test | Statistic method | *p* value | *t* value |
| --- | --- | --- | --- | --- | --- | --- |
| 2A | n = 12 M  n = 12 F | passed | passed | two-tailed *t*-test | *p* = 0.606 | *t* = 0.524 |
|  |  |  |  | two-tailed M-W *U* test | *p* = 0.799 | *U =* 67.000 |
| 2B (5 min) | n = 12 M  n = 12 F | passed | failed | two-tailed M-W *U* test | *p* = 0.160 | *U =* 47.500 |
| 2B (10 min) | n = 12 M  n = 12 F | failed |  | two-tailed M-W *U* test | *p* = 0.001 | *U =* 17.000 |
| 2B (15 min) | n = 12 M  n = 12 F | failed |  | two-tailed M-W *U* test | *p* < 0.001 | *U =* 9.000 |
| 2B (20 min) | n = 12 M  n = 12 F | failed |  | two-tailed M-W *U* test | *p* = 0.010 | *U =* 28.500 |
| 2B (25 min) | n = 12 M  n = 12 F | failed |  | two-tailed M-W *U* test | *p* = 0.378 | *U =* 56.000 |
| 2B (30 min) | n = 12 M  n = 12 F | failed |  | two-tailed M-W *U* test | *p* = 0.671 | *U =* 64.000 |
| 2C | n = 12 M  n = 12 F | passed | passed | two-tailed *t*-test | *p* = 0.262 | *t* = -1.151 |
|  |  |  |  | two-tailed M-W *U* test | *p* = 0.410 | *U =* 87.000 |
| 2D (5 min) | n = 12 M  n = 12 F | failed |  | two-tailed M-W *U* test | *p* = 0.033 | *U =* 35.500 |
| 2D (10 min) | n = 12 M  n = 12 F | failed |  | two-tailed M-W *U* test | *p* = 0.060 | *U =* 39.500 |
| 2D (15 min) | n = 12 M  n = 12 F | failed |  | two-tailed M-W *U* test | *p* = 0.178 | *U =* 48.000 |
| 2D (20 min) | n = 12 M  n = 12 F | failed |  | two-tailed M-W *U* test | *p* = 1.000 | *U =* 72.500 |
| 2D (25 min) | n = 12 M  n = 12 F | failed |  | two-tailed M-W *U* test | *p* = 0.977 | *U =* 71.500 |
| 2D (30 min) | n = 12 M  n = 12 F | failed |  | two-tailed M-W *U* test | *p* = 0.178 | *U =* 48.000 |
| 2E | n = 12 M  n = 12 F | failed |  | two-tailed M-W *U* test | *p* = 0.347 | *U =* 55.000 |
| 2F (5 min) | n = 12 M  n = 12 F | failed |  | two-tailed M-W *U* test | *p* = 0.143 | *U =* 46.500 |
| 2F (10 min) | n = 12 M  n = 12 F | failed |  | two-tailed M-W *U* test | *p* = 0.002 | *U =* 20.000 |
| 2F (15 min) | n = 12 M  n = 12 F | failed |  | two-tailed M-W *U* test | *p* = 0.347 | *U =* 55.500 |
| 2F (20 min) | n = 12 M  n = 12 F | failed |  | two-tailed M-W *U* test | *p* = 0.128 | *U =* 45.500 |
| 2F (25 min) | n = 12 M  n = 12 F | failed |  | two-tailed M-W *U* test | *p* = 0.089 | *U =* 42.000 |
| 2F (30 min) | n = 12 M  n = 12 F | failed |  | two-tailed M-W *U* test | *p* = 0.514 | *U =* 60.000 |
| 2G | n = 12 M  n = 12 F | passed | passed | two-tailed *t*-test | *p* = 0.251 | *t* = 1.179 |
|  |  |  |  | two-tailed M-W *U* test | *p* = 0.478 | *U =* 59.000 |
| 2H (5 min) | n = 12 M  n = 12 F | failed |  | two-tailed M-W *U* test | *p* = 0. 178 | *U =* 96.000 |
| 2H (10 min) | n = 12 M  n = 12 F | failed |  | two-tailed M-W *U* test | *p* = 0. 443 | *U =* 58.000 |
| 2H (15 min) | n = 12 M  n = 12 F | failed |  | two-tailed M-W *U* test | *p* = 0. 068 | *U =* 40.500 |
| 2H (20 min) | n = 12 M  n = 12 F | failed |  | two-tailed M-W *U* test | *p* = 0.775 | *U =* 78.000 |
| 2H (25 min) | n = 12 M  n = 12 F | failed |  | two-tailed M-W *U* test | *p* = 0.671 | *U =* 64.500 |
| 2H (30 min) | n = 12 M  n = 12 F | failed |  | two-tailed M-W *U* test | *p* = 0.630 | *U =* 80.500 |
| 2I | n = 12 M  n = 12 F | failed |  | two-tailed M-W *U* test | *p* = 0.525 | *U =* 55.500 |
| 2J (5min) | n = 12 M  n = 11 F | failed |  | two-tailed M-W *U* test | *p* = 0.786 | *U =* 70.500 |
| 2J (10min) | n = 12 M  n = 11 F | failed |  | two-tailed M-W *U* test | *p* = 0.151 | *U =* 42.500 |
| 2J (15min) | n = 12 M  n = 11 F | failed |  | two-tailed M-W *U* test | *p* = 0.169 | *U =* 43.000 |
| 2J (20min) | n = 12 M  n = 11 F | failed |  | two-tailed M-W *U* test | *p* = 0.786 | *U =* 61.500 |
| 2J (25min) | n = 12 M  n = 11 F | failed |  | two-tailed M-W *U* test | *p* = 0.928 | *U =* 67.500 |
| 2J (30min) | n = 12 M  n = 11 F | failed |  | two-tailed M-W *U* test | *p* = 0.487 | *U =* 54.500 |
| 2K | n = 12 M  n = 11 F | failed |  | two-tailed M-W *U* test | *p* = 0.487 | *U =* 54.500 |
| 2L (5min) | n = 12 M  n = 11 F | failed |  | two-tailed M-W *U* test | *p* = 0.928 | *U =* 68.000 |
| 2L (10min) | n = 12 M  n = 11 F | failed |  | two-tailed M-W *U* test | *p* = 0.134 | *U =* 91.000 |
| 2L (15min) | n = 12 M  n = 11 F | failed |  | two-tailed M-W *U* test | *p* = 0.695 | *U =* 59.500 |
| 2L (20min) | n = 12 M  n = 11 F | failed |  | two-tailed M-W *U* test | *p* = 0.740 | *U =* 72.000 |
| 2L (25min) | n = 12 M  n = 11 F | failed |  | two-tailed M-W *U* test | *p* = 0.740 | *U =* 60.500 |
| 2L (30min) | n = 12 M  n = 11 F | failed |  | two-tailed M-W *U* test | *p* = 0.740 | *U =* 60.500 |
| 2M | n = 12 M  n = 11 F | failed |  | two-tailed M-W *U* test | *p* = 0.019 | *U =* 104.000 |
| 2N (5min) | n = 12 M  n = 11 F | failed |  | two-tailed M-W *U* test | *p* = 0.316 | *U =* 49.000 |
| 2N (10min) | n = 12 M  n = 11 F | passed | passed | two-tailed *t*-test | *p* = 0.710 | *t* = 0.377 |
|  |  |  |  | two-tailed M-W *U* test | *p* = 0.740 | *U =* 60.500 |
| 2N (15min) | n = 12 M  n = 11 F | failed |  | two-tailed M-W *U* test | *p* = 0.413 | *U =* 80.000 |
| 2N (20min) | n = 12 M  n = 11 F | failed |  | two-tailed M-W *U* test | *p* = 0.379 | *U =* 80.500 |
| 2N (25min) | n = 12 M  n = 11 F | failed |  | two-tailed M-W *U* test | *p* = 0.651 | *U =* 58.500 |
| 3A | n = 12 M  n = 12 F | passed | failed | two-tailed M-W *U* test | *p* < 0.001 | *U =* 0.000 |
| 3B | n = 12 M  n = 12 F | failed |  | two-tailed M-W *U* test | *p* = 0.001 | *U =* 18.000 |
| 3C | n = 12 M  n = 12 F | passed | failed | two-tailed M-W *U* test | *p* = 0.004 | *U =* 23.000 |
| 3D | n = 12 M  n = 12 F | passed | failed | two-tailed M-W *U* test | *p* = 0.002 | *U =* 20.500 |
| 3E | n = 12 M  n = 12 F | failed |  | two-tailed M-W *U* test | *p* = 0.001 | *U =* 18.500 |
| 3F | n = 12 M  n = 12 F | failed |  | two-tailed M-W *U* test | *p* = 0.005 | *U =* 24.500 |
| 3G | n = 12 M  n = 12 F | failed |  | two-tailed M-W *U* test | *p* = 0.590 | *U =* 62.000 |
| 3H | n = 12 M  n = 12 F | failed |  | two-tailed M-W *U* test | *p* = 0.630 | *U =* 63.000 |
| 3I | n = 12 M  n = 11 F | passed | passed | two-tailed *t*-test | *p* = 0.388 | *t* = 0.881 |
|  |  |  |  | two-tailed M-W *U* test | *p* = 0.566 | *U =* 56.000 |
| 3J | n = 12 M  n = 11 F | passed | passed | two-tailed *t*-test | *p* = 0.266 | *t* = 1.142 |
|  |  |  |  | two-tailed M-W *U* test | *p* = 0.347 | *U =* 50.000 |
| 3K | n = 12 M  n = 11 F | failed |  | two-tailed M-W *U* test | *p* = 0.880 | *U =* 69.000 |
| 3L | n = 12 M  n = 11 F | passed | passed | two-tailed *t*-test | *p* = 0.259 | *t* = -1.161 |
|  |  |  |  | two-tailed M-W *U* test | *p* = 0.449 | *U =* 79.000 |
| 3M | n = 12 M  n = 11 F | passed | passed | two-tailed *t*-test | *p* = 0.644 | *t* = 0.469 |
|  |  |  |  | two-tailed M-W *U* test | *p* = 1.000 | *U =* 67.000 |
| 3N | n = 12 M  n = 11 F | passed | passed | two-tailed *t*-test | *p* = 0.702 | *t* = -0.388 |
|  |  |  |  | two-tailed M-W *U* test | *p* = 0.880 | *U =* 69.000 |
| 4 COnaive-CDnaive Baseline (0.16 g) | n = 12 M  n = 12 F | failed |  | two-tailed M-W *U* test | *p* = 0.755 | *U =* 66.000 |
| 4 COnaive-CDnaive Baseline (0.40 g) | n = 12 M  n = 12 F | failed |  | two-tailed M-W *U* test | *p* = 1.000 | *U =* 72.500 |
| 4 COnaive-CDnaive Baseline (0.60 g) | n = 12 M  n = 12 F | failed |  | two-tailed M-W *U* test | *p* = 0.755 | *U =* 77.500 |
| 4 COnaive-CDnaive Baseline (1.00 g) | n = 12 M  n = 12 F | failed |  | two-tailed M-W *U* test | *p* = 1.000 | *U =* 72.500 |
| 4 COnaive-CDnaive Baseline (1.40 g) | n = 12 M  n = 12 F | failed |  | two-tailed M-W *U* test | *p* = 0.219 | *U =* 94.000 |
| 4A (0.16 g) | n = 12 M  n = 12 F | failed |  | two-tailed M-W *U* test | *p* = 0.755 | *U =* 78.000 |
| 4A (0.40 g) | n = 12 M  n = 12 F | failed |  | two-tailed M-W *U* test | *p* = 0.347 | *U =* 88.500 |
| 4A (0.60 g) | n = 12 M  n = 12 F | failed |  | two-tailed M-W *U* test | *p* = 0.551 | *U =* 61.000 |
| 4A (1.00 g) | n = 12 M  n = 12 F | failed |  | two-tailed M-W *U* test | *p* = 0.242 | *U =* 92.500 |
| 4A (1.40 g) | n = 12 M  n = 12 F | failed |  | two-tailed M-W *U* test | *p* = 0.799 | *U =* 76.500 |
| 4B (0.16 g) | n = 12 M  n = 12 F | failed |  | two-tailed M-W *U* test | *p* = 1.000 | *U =* 72.000 |
| 4B (0.40 g) | n = 12 M  n = 12 F | failed |  | two-tailed M-W *U* test | *p* = 0.242 | *U =* 51.500 |
| 4B (0.60 g) | n = 12 M  n = 12 F | failed |  | two-tailed M-W *U* test | *p* = 0. 755 | *U =* 66.000 |
| 4B (1.00 g) | n = 12 M  n = 12 F | failed |  | two-tailed M-W *U* test | *p* = 0.755 | *U =* 78.000 |
| 4B (1.40 g) | n = 12 M  n = 12 F | failed |  | two-tailed M-W *U* test | *p* = 1.000 | *U =* 72.000 |
| 4C (0.16 g) | n = 12 M  n = 12 F | failed |  | two-tailed M-W *U* test | *p* = 0.514 | *U =* 84.000 |
| 4C (0.40 g) | n = 12 M  n = 12 F | failed |  | two-tailed M-W *U* test | *p* = 0.551 | *U =* 83.000 |
| 4C (0.60 g) | n = 12 M  n = 12 F | failed |  | two-tailed M-W *U* test | *p* = 0.347 | *U =* 88.500 |
| 4C (1.00 g) | n = 12 M  n = 12 F | failed |  | two-tailed M-W *U* test | *p* = 0.219 | *U =* 50.000 |
| 4C (1.40 g) | n = 12 M  n = 12 F | failed |  | two-tailed M-W *U* test | *p* = 0.378 | *U =* 56.000 |
| 4D (0.16 g) | n = 12 M  n = 12 F | failed |  | two-tailed M-W *U* test | *p* = 1.000 | *U =* 72.000 |
| 4D (0.40 g) | n = 12 M  n = 12 F | failed |  | two-tailed M-W *U* test | *p* = 0.799 | *U =* 67.000 |
| 4D (0.60 g) | n = 12 M  n = 12 F | failed |  | two-tailed M-W *U* test | *p* = 0.551 | *U =* 83.000 |
| 4D (1.00 g) | n = 12 M  n = 12 F | failed |  | two-tailed M-W *U* test | *p* = 0.755 | *U =* 77.500 |
| 4D (1.40 g) | n = 12 M  n = 12 F | failed |  | two-tailed M-W *U* test | *p* = 0.755 | *U =* 78.000 |
| 4E (0.16 g) | n = 12 M  n = 12 F | failed |  | two-tailed M-W *U* test | *p* = 0.755 | *U =* 66.000 |
| 4E (0.40 g) | n = 12 M  n = 12 F | failed |  | two-tailed M-W *U* test | *p* = 0.242 | *U =* 51.000 |
| 4E (0.60 g) | n = 12 M  n = 12 F | failed |  | two-tailed M-W *U* test | *p* = 0.755 | *U =* 66.500 |
| 4E (1.00 g) | n = 12 M  n = 12 F | failed |  | two-tailed M-W *U* test | *p* = 0.514 | *U =* 84.000 |
| 4E (1.40 g) | n = 12 M  n = 12 F | failed |  | two-tailed M-W *U* test | *p* = 0.755 | *U =* 66.500 |
| 4F (0.16 g) | n = 12 M  n = 12 F | failed |  | two-tailed M-W *U* test | *p* = 0.178 | *U =* 96.000 |
| 4F (0.40 g) | n = 12 M  n = 12 F | failed |  | two-tailed M-W *U* test | *p* = 0.799 | *U =* 67.000 |
| 4F (0.60 g) | n = 12 M  n = 12 F | failed |  | two-tailed M-W *U* test | *p* = 0.515 | *U =* 83.500 |
| 4F (1.00 g) | n = 12 M  n = 12 F | failed |  | two-tailed M-W *U* test | *p* = 0.932 | *U =* 70.500 |
| 4F (1.40 g) | n = 12 M  n = 12 F | failed |  | two-tailed M-W *U* test | *p* = 0.755 | *U =* 66.000 |
| 4 COnaive-CDpain Baseline (0.16 g) | n = 12 M  n = 12 F | failed |  | two-tailed M-W *U* test | *p* = 0.755 | *U =* 78.000 |
| 4 COnaive-CDpain Baseline (0.40 g) | n = 12 M  n = 12 F | failed |  | two-tailed M-W *U* test | *p* = 0.514 | *U =* 84.000 |
| 4 COnaive-CDpain Baseline (0.60 g) | n = 12 M  n = 12 F | failed |  | two-tailed M-W *U* test | *p* = 0.443 | *U =* 58.500 |
| 4 COnaive-CDpain Baseline (1.00 g) | n = 12 M  n = 12 F | failed |  | two-tailed M-W *U* test | *p* = 0.443 | *U =* 58.000 |
| 4 COnaive-CDpain Baseline (1.40 g) | n = 12 M  n = 12 F | failed |  | two-tailed M-W *U* test | *p* = 0.410 | *U =* 57.000 |
| 4G (0.16 g) | n = 12 M  n = 12 F | failed |  | two-tailed M-W *U* test | *p* = 0.242 | *U =* 51.000 |
| 4G (0.40 g) | n = 12 M  n = 12 F | failed |  | two-tailed M-W *U* test | *p* = 0.178 | *U =* 96.000 |
| 4G (0.60 g) | n = 12 M  n = 12 F | failed |  | two-tailed M-W *U* test | *p* = 0.630 | *U =* 63.000 |
| 4G (1.00 g) | n = 12 M  n = 12 F | failed |  | two-tailed M-W *U* test | *p* = 0.198 | *U =* 49.500 |
| 4G (1.40 g) | n = 12 M  n = 12 F | failed |  | two-tailed M-W *U* test | *p* = 0.755 | *U =* 78.000 |
| 4H (0.16 g) | n = 12 M  n = 12 F | failed |  | two-tailed M-W *U* test | *p* = 0.101 | *U =* 43.500 |
| 4H (0.40 g) | n = 12 M  n = 12 F | failed |  | two-tailed M-W *U* test | *p* = 0.319 | *U =* 54.000 |
| 4H (0.60 g) | n = 12 M  n = 12 F | failed |  | two-tailed M-W *U* test | *p* = 0.347 | *U =* 55.500 |
| 4H (1.00 g) | n = 12 M  n = 12 F | failed |  | two-tailed M-W *U* test | *p* = 0.443 | *U =* 58.000 |
| 4H (1.40 g) | n = 12 M  n = 12 F | failed |  | two-tailed M-W *U* test | *p* = 0.319 | *U =* 54.000 |
| 4I (0.16 g) | n = 12 M  n = 12 F | failed |  | two-tailed M-W *U* test | *p* = 0.219 | *U =* 50.000 |
| 4I (0.40 g) | n = 12 M  n = 12 F | failed |  | two-tailed M-W *U* test | *p* = 0.410 | *U =* 57.500 |
| 4I (0.60 g) | n = 12 M  n = 12 F | failed |  | two-tailed M-W *U* test | *p* = 0.143 | *U =* 46.500 |
| 4I (1.00 g) | n = 12 M  n = 12 F | failed |  | two-tailed M-W *U* test | *p* = 0.143 | *U =* 46.500 |
| 4I (1.40 g) | n = 12 M  n = 12 F | failed |  | two-tailed M-W *U* test | *p* = 0.932 | *U =* 74.000 |
| 4J (0.16 g) | n = 12 M  n = 12 F | failed |  | two-tailed M-W *U* test | *p* = 0.101 | *U =* 43.500 |
| 4J (0.40 g) | n = 12 M  n = 12 F | failed |  | two-tailed M-W *U* test | *p* = 0.242 | *U =* 51.000 |
| 4J (0.60 g) | n = 12 M  n = 12 F | failed |  | two-tailed M-W *U* test | *p* = 0.630 | *U =* 63.000 |
| 4J (1.00 g) | n = 12 M  n = 12 F | failed |  | two-tailed M-W *U* test | *p* = 0.143 | *U =* 46.500 |
| 4J (1.40 g) | n = 12 M  n = 12 F | failed |  | two-tailed M-W *U* test | *p* = 0.932 | *U =* 74.000 |
| 4K (0.16 g) | n = 12 M  n = 12 F | failed |  | two-tailed M-W *U* test | *p* = 0.052 | *U =* 38.500 |
| 4K (0.40 g) | n = 12 M  n = 12 F | failed |  | two-tailed M-W *U* test | *p* = 0.128 | *U =* 45.000 |
| 4K (0.60 g) | n = 12 M  n = 12 F | failed |  | two-tailed M-W *U* test | *p* = 0.002 | *U =* 21.000 |
| 4K (1.00 g) | n = 12 M  n = 12 F | failed |  | two-tailed M-W *U* test | *p* = 0.014 | *U =* 30.000 |
| 4K (1.40 g) | n = 12 M  n = 12 F | failed |  | two-tailed M-W *U* test | *p* = 0.114 | *U =* 44.500 |
| 4L (0.16 g) | n = 12 M  n = 12 F | failed |  | two-tailed M-W *U* test | *p* = 0.089 | *U =* 42.500 |
| 4L (0.40 g) | n = 12 M  n = 12 F | failed |  | two-tailed M-W *U* test | *p* = 0.143 | *U =* 46.000 |
| 4L (0.60 g) | n = 12 M  n = 12 F | failed |  | two-tailed M-W *U* test | *p* = 0.266 | *U =* 52.000 |
| 4L (1.00 g) | n = 12 M  n = 12 F | failed |  | two-tailed M-W *U* test | *p* = 0.219 | *U =* 50.000 |
| 4L (1.40 g) | n = 12 M  n = 12 F | failed |  | two-tailed M-W *U* test | *p* = 0.078 | *U =* 41.500 |
| 5A (BL) | n = 12 M  n = 12 F | passed | passed | two-tailed *t*-test | *p* = 0.436 | *t* = 0.793 |
|  |  |  |  | two-tailed M-W *U* test | *p* = 0.478 | *U =* 59.000 |
| 5A (PDSI) | n = 12 M  n = 12 F | passed | passed | two-tailed *t*-test | *p* = 0.450 | *t* = 0.769 |
|  |  |  |  | two-tailed M-W *U* test | *p* = 0.977 | *U =* 71.500 |
| 5A (60 min) | n = 12 M  n = 12 F | failed |  | two-tailed M-W *U* test | *p* = 0.671 | *U =* 80.000 |
| 5A (120 min) | n = 12 M  n = 12 F | failed |  | two-tailed M-W *U* test | *p* = 0.630 | *U =* 63.500 |
| 5A (180 min) | n = 12 M  n = 12 F | failed |  | two-tailed M-W *U* test | *p* = 0.977 | *U =* 71.000 |
| 5A (240 min) | n = 12 M  n = 12 F | passed | passed | two-tailed *t*-test | *p* = 0.353 | *t* = -0.950 |
|  |  |  |  | two-tailed M-W *U* test | *p* = 0.319 | *U =* 89.500 |
| 5A (300 min) | n = 12 M  n = 12 F | failed |  | two-tailed M-W *U* test | *p* = 0.755 | *U =* 78.000 |
| 5B (BL) | n = 12 M  n = 12 F | passed | failed | two-tailed M-W *U* test | *p* = 0.843 | *U =* 76.000 |
| 5B (PDSI) | n = 12 M  n = 12 F | failed |  | two-tailed M-W *U* test | *p* = 0.755 | *U =* 66.000 |
| 5B (60 min) | n = 12 M  n = 12 F | failed |  | two-tailed M-W *U* test | *p* = 0.114 | *U =* 100.000 |
| 5B (120 min) | n = 12 M  n = 12 F | passed | failed | two-tailed M-W *U* test | *p* = 0.443 | *U =* 86.000 |
| 5B (180 min) | n = 12 M  n = 12 F | failed |  | two-tailed M-W *U* test | *p* = 0.291 | *U =* 91.000 |
| 5B (240 min) | n = 12 M  n = 12 F | passed | failed | two-tailed M-W *U* test | *p* = 0.003 | *U =* 122.000 |
| 5B (300 min) | n = 12 M  n = 12 F | passed | failed | two-tailed M-W *U* test | *p* = 0.060 | *U =* 105.000 |
| 5C (BL) | n = 8 M  n = 9 F | failed |  | two-tailed M-W *U* test | *p* = 1.000 | *U =* 37.000 |
| 5C (PDSI) | n = 8 M  n = 9 F | failed |  | two-tailed M-W *U* test | *p* = 0.370 | *U =* 46.000 |
| 5C (30min) | n = 8 M  n = 9 F | failed |  | two-tailed M-W *U* test | *p* = 0.370 | *U =* 46.000 |
| 5C (60min) | n = 8 M  n = 9 F | failed |  | two-tailed M-W *U* test | *p* = 1.000 | *U =* 36.000 |
| 5C (120min) | n = 8 M  n = 9 F | failed |  | two-tailed M-W *U* test | *p* = 0.423 | *U =* 45.000 |
| 5C (180min) | n = 8 M  n = 9 F | failed |  | two-tailed M-W *U* test | *p* = 0.888 | *U =* 38.000 |
| 5C (240min) | n = 8 M  n = 9 F | passed | passed | two-tailed *t*-test | *p* = 0.694 | *t* = 0.401 |
|  |  |  |  | two-tailed M-W *U* test | *p* = 0.888 | *U =* 34.500 |
| 5C (300min) | n = 8 M  n = 9 F | failed |  | two-tailed M-W *U* test | *p* = 0.481 | *U =* 44.000 |
| 5D (BL) | n = 8 M  n = 12 F | failed |  | two-tailed M-W *U* test | *p* = 1.000 | *U =* 48.000 |
| 5D (PDSI) | n = 8 M  n = 12 F | failed |  | two-tailed M-W *U* test | *p* = 0.734 | *U =* 43.500 |
| 5D (30min) | n = 8 M  n = 12 F | failed |  | two-tailed M-W *U* test | *p* = 0.851 | *U =* 45.500 |
| 5D (60min) | n = 8 M  n = 12 F | passed | passed | two-tailed *t*-test | *p* = 0.511 | *t* = -0.671 |
|  |  |  |  | two-tailed M-W *U* test | *p* = 0.571 | *U =* 55.500 |
| 5D (120min) | n = 8 M  n = 12 F | passed | passed | two-tailed *t*-test | *p* = 0.444 | *t* = -0.783 |
|  |  |  |  | two-tailed M-W *U* test | *p* = 0.571 | *U =* 56.000 |
| 5D (180min) | n = 8 M  n = 12 F | failed |  | two-tailed M-W *U* test | *p* = 0.208 | *U =* 65.000 |
| 5E (BL) | n = 8 M  n = 9 F | failed |  | two-tailed M-W *U* test | *p* = 0.541 | *U =* 43.000 |
| 5E (PDSI) | n = 8 M  n = 9 F | failed |  | two-tailed M-W *U* test | *p* = 1.000 | *U =* 36.500 |
| 5E (30min) | n = 8 M  n = 9 F | failed |  | two-tailed M-W *U* test | *p* =1.000 | *U =* 36.000 |
| 5E (60min) | n = 8 M  n = 9 F | failed |  | two-tailed M-W *U* test | *p* = 0.541 | *U =* 43.000 |
| 5E (120min) | n = 8 M  n = 9 F | failed |  | two-tailed M-W *U* test | *p* = 0.606 | *U =* 41.500 |
| 5E (180min) | n = 8 M  n = 9 F | failed |  | two-tailed M-W *U* test | *p* = 0.673 | *U =* 31.500 |
| 5E (240min) | n = 8 M  n = 9 F | failed |  | two-tailed M-W *U* test | *p* = 0.277 | *U =* 24.000 |
| 5E (300min) | n = 8 M  n = 9 F | failed |  | two-tailed M-W *U* test | *p* = 0.743 | *U =* 32.000 |
| 5F (BL) | n = 8 M  n = 12 F | failed |  | two-tailed M-W *U* test | *p* = 0.473 | *U =* 58.000 |
| 5F (PDSI) | n = 8 M  n = 12 F | passed | passed | two-tailed *t*-test | *p* = 0.686 | *t* = -0.410 |
|  |  |  |  | two-tailed M-W *U* test | *p* = 0.851 | *U =* 45.500 |
| 5F (30min) | n = 8 M  n = 12 F | failed |  | two-tailed M-W *U* test | *p* = 0.792 | *U =* 44.000 |
| 5F (60min) | n = 8 M  n = 12 F | passed | passed | two-tailed *t*-test | *p* = 0.837 | *t* = -0.209 |
|  |  |  |  | two-tailed M-W *U* test | *p* = 0.792 | *U =* 51.500 |
| 5F (120min) | n = 8 M  n = 12 F | passed | passed | two-tailed *t*-test | *p* = 0.824 | *t* = -0.225 |
|  |  |  |  | two-tailed M-W *U* test | *p* = 0.851 | *U =* 50.500 |
| 5F (180min) | n = 8 M  n = 12 F | passed | failed | two-tailed M-W *U* test | *p* = 0.270 | *U =* 63.000 |
| 5F (240min) | n = 8 M  n = 12 F | passed | passed | two-tailed *t*-test | *p* = 0.839 | *t* = -0.207 |
|  |  |  |  | two-tailed M-W *U* test | *p* = 0.734 | *U =* 53.000 |
| 5F (300min) | n = 8 M  n = 12 F | failed |  | two-tailed M-W *U* test | *p* = 0.678 | *U =* 54.000 |

Notes: The values of both parametric and non-parametric statistical analysis methods were presented if both normality test and equal variance test for samples passed, while, only results of non-parametric statistical analysis method were presented if either the normality test or equal variance test failed. *p* < 0.05 was considered as statistically significant. M, male; F, female; *t*-test, two-sample *t*-test; M-W *U* test, Mann-Whitney *U* test.
