## Supplemental Table SII for "Empathic contagious pain and consolation in laboratory rodents: species and sex comparisons"

Table SII Time effects of empathic consoling and empathic contagion of pain in mice and rats of both sexes

| Figure number | Number of animals | Statistic method 1 | Between-sample effects | Within-time effects | Statistic method 2 | *p* values |
| --- | --- | --- | --- | --- | --- | --- |
| 2B | n = 12 M  n = 12 F | Two-way ANOVA RM with Bonferroni post hoc correction | *p* < 0.001  *F_A_* = *F*_0.05, (1,22)_ = 57.510 | Mauchly's Test of Sphericity  𝜒^2^ = 32.600  𝜈 = 14  *p* = 0.003 (failed)  then,  Greenhouse-Geisser:  *p* < 0.001  *F_B_* = *F*_0.05, (3.709,81.596)_ = 8.952  Greenhouse-Geisser:  *p* = 0.231  *F_AB_* = *F*_0.05, (3.709,81.596)_ = 1.441 | Wilcoxon Signed Rank Test  *𝛂’* = 0.01 | M (5 min vs 10 min): *p* = 0.202  M (10 min vs 15 min): *p* = 0.423  M (15 min vs 20 min): *p* = 0.937  M (20 min vs 25 min): *p* = 0.025  M (25 min vs 30 min): *p* = 0.734  F (5 min vs 10 min): *p* = 0.003  F (10 min vs 15 min): *p* = 0.167  F (15 min vs 20 min): *p* = 0.125  F (20 min vs 25 min): *p* = 0.042  F (25 min vs 30 min): *p* = 0.072 |
| 2D | n = 12 M  n = 12 F | Two-way ANOVA RM with Bonferroni post hoc correction | *p* = 0.005  *F_A_* = *F*_0.05, (1,22)_ = 9.816 | Mauchly's Test of Sphericity  𝜒^2^ = 54.330  𝜈 = 14  *p* < 0.001 (failed)  then,  Greenhouse-Geisser: *p* = 0.015  *F_B_* = *F*_0.05, (2.981,65.591)_ = 3.778  Greenhouse-Geisser: *p* = 0.204  *F_AB_* = *F*_0.05, (2.981,65.591)_ = 1.574 | Wilcoxon Signed Rank Test  *𝛂’* = 0.01 | M (5 min vs 10 min): *p*= 0.183  M (10 min vs 15 min): *p* = 0.322  M (15 min vs 20 min): *p* = 0.833  M (20 min vs 25 min): *p* = 0.276  M (25 min vs 30 min): *p* = 0.257  F (5 min vs 10 min): *p* = 0.042  F (10 min vs 15 min): *p* = 0.317  F (15 min vs 20 min): *p* = 0.066  F (20 min vs 25 min): *p* = 0.109  F (25 min vs 30 min): *p* = 0.317 |
| 2F | n = 12 M  n = 12 F | Two-way ANOVA RM with Bonferroni post hoc correction | *p* = 0.002  *F_A_* = *F*_0.05, (1,22)_ = 13.069 | Mauchly's Test of Sphericity  𝜒^2^ = 79.733  𝜈 = 14  *p* < 0.001 (failed)  then,  Greenhouse-Geisser: *p* = 0.017  *F_B_* = *F*_0.05, (2.127,46.792)_ = 4.348  Greenhouse-Geisser: *p* = 0.192  *F_AB_* = *F*_0.05, (2.127,46.792)_ = 1.703 | Wilcoxon Signed Rank Test  *𝛂’* = 0.01 | M (5 min vs 10 min): *p*= 0.159  M (10 min vs 15 min): *p* = 0.050  M (15 min vs 20 min): *p* = 0.128  M (20 min vs 25 min): *p* = 0.176  M (25 min vs 30 min): *p* = 0.236  F (5 min vs 10 min): *p* = 0.017  F (10 min vs 15 min): *p* = 0.655  F (15 min vs 20 min): *p* = 0.655  F (20 min vs 25 min): *p* = 0.317  F (25 min vs 30 min): *p* = 1.000 |
| 2H | n = 12 M  n = 12 F | Two-way ANOVA RM with Bonferroni post hoc correction | *p* = 0.795  *F_A_* = *F*_0.05, (1,22)_ = 0.070 | Mauchly's Test of Sphericity  𝜒^2^ = 24.910  𝜈 = 14  *p* = 0.037 (failed)  then,  Greenhouse-Geisser: *p* = 0.027  *F_B_* = *F*_0.05, (3.525,77.547)_ = 3.048  Greenhouse-Geisser: *p* = 0.166  *F_AB_* = *F*_0.05, (3.525,77.547)_ = 1.698 | Wilcoxon Signed Rank Test  *𝛂’* = 0.01 | M (5 min vs 10 min): *p* = 0.066  M (10 min vs 15 min): *p* = 0.136  M (15 min vs 20 min): *p* = 0.114  M (20 min vs 25 min): *p* = 0.058  M (25 min vs 30 min): *p* = 0.498  F (5 min vs 10 min): *p* = 1.000  F (10 min vs 15 min): *p* = 0.866  F (15 min vs 20 min): *p* = 0.575  F (20 min vs 25 min): *p* = 0.086  F (25 min vs 30 min): *p* = 0.075 |
| 2J | n = 12 M  n = 11 F | Two-way ANOVA RM with Bonferroni post hoc correction | *p* = 0.388  *F_A_* = *F*_0.05, (1,21)_ = 0.776 | Mauchly's Test of Sphericity  𝜒^2^ = 38.388  𝜈 = 14  *p* < 0.001 (failed)  then,  Greenhouse-Geisser: *p* = 0.002  *F_B_* = *F*_0.05, (2.926,61.450)_ = 5.650  Greenhouse-Geisser: *p* = 0.618  *F_AB_*= *F*_0.05, (2.926,61.450)_ = 0.592 | Wilcoxon Signed Rank Test  *𝛂’* = 0.01 | M (5 min vs 10 min): *p*= 0.556  M (10 min vs 15 min): *p* = 0.126  M (15 min vs 20 min): *p* = 0.102  M (20 min vs 25 min): *p* = 0.594  M (25 min vs 30 min): *p* = 0.673  F (5 min vs 10 min): *p* = 0.003  F (10 min vs 15 min): *p* = 0.063  F (15 min vs 20 min): *p* = 0.528  F (20 min vs 25 min): *p* = 0.916  F (25 min vs 30 min): *p* = 0.574 |
| 2L | n = 12 M  n = 11 F | Two-way ANOVA RM with Bonferroni post hoc correction | *p* = 0.484  *F_A_* = *F*_0.05, (1,21)_ = 0.507 | Mauchly's Test of Sphericity  𝜒^2^ = 161.951  𝜈 = 14  *p* < 0.001 (failed)  then,  Greenhouse-Geisser: *p* = 0.001  *F_B_* = *F*_0.05, (1.614,33.898)_ = 10.406  Greenhouse-Geisser: *p* = 0.469  *F_AB_* = *F*_0.05, (1.614,33.898)_ = 0.714 | Wilcoxon Signed Rank Test  *𝛂’* = 0.01 | M (5 min vs 10 min): *p*= 0.021  M (10 min vs 15 min): *p* = 0.416  M (15 min vs 20 min): *p* = 0.109  M (20 min vs 25 min): *p* = 0.317  M (25 min vs 30 min): *p* = 0.655  F (5 min vs 10 min): *p* = 0.201  F (10 min vs 15 min): *p* = 0.034  F (15 min vs 20 min): *p* = 0.564  F (20 min vs 25 min): *p* = 0.317  F (25 min vs 30 min): *p* = 1.000 |
| 2N | n = 12 M  n = 11 F | Two-way ANOVA RM with Bonferroni post hoc correction | *p* = 0.536  *F_A_* = *F*_0.05, (1,21)_ = 0.397 | Mauchly's Test of Sphericity  𝜒^2^ = 38.536  𝜈 = 14  *p* < 0.001 (failed)  then,  Greenhouse-Geisser: *p* = 0.013  *F_B_* = *F*_0.05, (3.025,63.523)_ = 3.850  Greenhouse-Geisser: *p* = 0.485  *F_AB_* = *F*_0.05, (3.025,63.523)_ = 0.827 | Wilcoxon Signed Rank Test  *𝛂’* = 0.01 | M (5 min vs 10 min): *p* = 0.374  M (10 min vs 15 min): *p* = 0.182  M (15 min vs 20 min): *p* = 0.594  M (20 min vs 25 min): *p* = 0.624  M (25 min vs 30 min): *p* = 0.028  F (5 min vs 10 min): *p* = 0.023  F (10 min vs 15 min): *p* = 0.838  F (15 min vs 20 min): *p* = 0.594  F (20 min vs 25 min): *p* = 0.169  F (25 min vs 30 min): *p* = 0.028 |
| 5A | n = 12 M  n = 12 F | Two-way ANOVA RM with Bonferroni post hoc correction | *p* = 0.737  *F_A_* = *F*_0.05, (1,22)_ = 0.116 | Mauchly's Test of Sphericity  𝜒^2^ = 34.014  𝜈 = 20  *p* = 0.028 (failed)  then,  Greenhouse-Geisser:  *p* = 0.251  *F_B_* = *F*_0.05, (3.501,77.017)_ = 1.384 *p* = 0.790  *F_AB_* = *F*_0.05, (3.501,77.017)_ = 0.390 | Wilcoxon Signed Rank Test  *𝛂’* = 0.008 | M (BL vs PDSI): *p* = 0.575  M (PDSI vs 60 min): *p* = 0.814  M (60 min vs 120 min): *p* = 0.190  M (120 min vs 180 min): *p* = 1.000  M (180 min vs 240 min): *p* = 0.859  M (240 min vs 300 min): *p* = 0.114  F (BL vs PDSI): *p* = 0.477  F (PDSI vs 60 min): *p* = 0.314  F (60 min vs 120 min): *p* = 0.594  F (120 min vs 180 min): *p* = 0.760  F (180 min vs 240 min): *p* = 0.173  F (240 min vs 300 min): *p* = 0.262 |
|  |  |  |  |  | Wilcoxon Signed Rank Test  *𝛂’* = 0.008 | M (BL vs PDSI): *p* = 0.575  M (BL vs 60 min): *p* = 0.139  M (BL vs 120 min): *p* = 0.308  M (BL vs 180 min): *p* = 0.790  M (BL vs 240 min): *p* = 0.433  M (BL vs 300 min): *p* = 0.646  F (BL vs PDSI): *p* = 0.477  F (BL vs 60 min): *p* = 0.959  F (BL vs 120 min): *p* = 0.689  F (BL vs 180 min): *p* = 0.508  F (BL vs 240 min): *p* = 0.530  F (BL vs 300 min): *p* = 0.477 |
| 5B | n = 12 M  n = 12 F | Two-way ANOVA RM with Bonferroni post hoc correction | *p* = 0.156  *F_A_* = *F*_0.05, (1,22)_ = 2.161 | Mauchly's Test of Sphericity  𝜒^2^ = 29.452  𝜈 = 20  *p*= 0.083 (passed)  then,  *p* < 0.001  *F_B_* = *F*_0.05, (6,132)_ = 24.029 *p* = 0.007  *F_AB_* = *F*_0.05, (6,132)_ = 3.131 | Wilcoxon Signed Rank Test  *𝛂’* = 0.008 | M (BL vs PDSI): *p* = 0.003  M (PDSI vs 60 min): *p* = 0.028  M (60 min vs 120 min): *p* = 0.208  M (120 min vs 180 min): *p* = 0.508  M (180 min vs 240 min): *p* = 0.285  M (240 min vs 300 min): *p* = 0.041  F (BL vs PDSI): *p* = 0.002  F (PDSI vs 60 min): *p* = 0.875  F (60 min vs 120 min): *p* = 0.514  F (120 min vs 180 min): *p* = 0.594  F (180 min vs 240 min): *p* = 0.013  F (240 min vs 300 min): *p* = 0.424 |
|  |  |  |  |  | Wilcoxon Signed Rank Test  *𝛂’* = 0.008 | M (BL vs PDSI): *p* = 0.003  M (BL vs 60 min): *p* = 0.002  M (BL vs 120 min): *p* = 0.002  M (BL vs 180 min): *p* = 0.003  M (BL vs 240 min): *p* = 0.002  M (BL vs 300 min): *p* = 0.004  F (BL vs PDSI): *p* = 0.002  F (BL vs 60 min): *p* = 0.003  F (BL vs 120 min): *p* = 0.002  F (BL vs 180 min): *p* = 0.002  F (BL vs 240 min): *p* = 0.009  F (BL vs 300 min): *p* = 0.003 |
| 5C | n = 8 M  n = 9 F | Two-way ANOVA RM with Bonferroni post hoc correction | *p* = 0.467  *F_A_* = *F*_0.05, (1,15)_ = 0.557 | Mauchly's Test of Sphericity  𝜒^2^ = 50.666  𝜈 = 27  *p* = 0.005 (failed)  then,  Greenhouse-Geisser:  *p* = 0.578  *F_B_* = *F*_0.05, (2.768,41.519)_ = 0.645 Greenhouse-Geisser:  *p* = 0.708  *F_AB_* = *F*_0.05, (2.768,41.519)_ = 0.443 | Wilcoxon Signed Rank Test  *𝛂’* = 0.008 | M (BL vs PDSI): *p* = 0.669  M (PDSI vs 30 min): *p* = 1.000  M (30 min vs 60 min): *p* = 0.285  M (60 min vs 120 min): *p* = 0.144  M (120 min vs 180 min): *p* = 0.180  M (180 min vs 240 min): *p* = 0.102  M (240 min vs 300 min): *p* = 0.785  F (BL vs PDSI): *p* = 0.859  F (PDSI vs 30 min): *p* = 0.892  F (30 min vs 60 min): *p* = 0.269  F (60 min vs 120 min): *p* = 0.785  F (120 min vs 180 min): *p* = 1.000  F (180 min vs 240 min): *p* = 0.863  F (240 min vs 300 min): *p* = 0.269 |
|  |  |  |  |  | Wilcoxon Signed Rank Test  *𝛂’* = 0.008 | M (BL vs PDSI): *p* = 0.669  M (BL vs 30 min): *p* = 0.260  M (BL vs 60 min): *p* = 0.888  M (BL vs 120 min): *p* = 0.260  M (BL vs 180 min):*p* = 0.670  M (BL vs 240 min): *p* = 1.000  M (BL vs 300 min): *p* = 1.000  F (BL vs PDSI): *p* = 0.859  F (BL vs 30 min): *p* = 0.767  F (BL vs 60 min): *p* = 0.593  F (BL vs 120 min): *p* = 0.593  F (BL vs 180 min): *p* = 0.513  F (BL vs 240 min): *p* = 0.859  F (BL vs 300 min): *p* = 0.311 |
| 5D | n = 8 M  n = 12 F | Two-way ANOVA RM with Bonferroni post hoc correction | *p* = 0.444  *F_A_* = *F*_0.05, (1,18)_ = 0.612 | Mauchly's Test of Sphericity  𝜒^2^ = 109.912  𝜈 = 27  *p* < 0.001 (failed)  then,  Greenhouse-Geisser:  *p* < 0.001  *F_B_* = *F*_0.05, (2.575,46.348)_ = 54.758 Greenhouse-Geisser:  *p* = 0.666  *F_AB_* = *F*_0.05, (2.575,46.348)_ = 0.484 | Wilcoxon Signed Rank Test  *𝛂’* = 0.008 | M (BL vs PDSI): *p* = 0.012  M (PDSI vs 30 min): *p* = 1.000  M (30 min vs 60 min): *p* = 0.317  M (60 min vs 120 min): *p* = 0.066  M (120 min vs 180 min): *p* = 0.180  M (180 min vs 240 min): *p* = 0.028  M (240 min vs 300 min): *p* = 0.018  F (BL vs PDSI): *p* = 0.002  F (PDSI vs 30 min): *p* = 0.180  F (30 min vs 60 min): *p* = 0.345  F (60 min vs 120 min): *p* = 0.043  F (120 min vs 180 min): *p* = 0.686  F (180 min vs 240 min): *p* = 0.407  F (240 min vs 300 min): *p* = 0.005 |
|  |  |  |  |  | Wilcoxon Signed Rank Test  *𝛂’* = 0.008 | M (BL vs PDSI): *p* = 0.012  M (BL vs 30 min): *p* = 0.012  M (BL vs 60 min): *p* = 0.012  M (BL vs 120 min): *p* = 0.012  M (BL vs 180 min):*p* = 0.012  M (BL vs 240 min): *p* = 0.011  M (BL vs 300 min): *p* = 1.000  F (BL vs PDSI): *p* = 0.002  F (BL vs 30 min): *p* = 0.002  F (BL vs 60 min): *p* = 0.002  F (BL vs 120 min): *p* = 0.002  F (BL vs 180 min): *p* = 0.002  F (BL vs 240 min): *p* = 0.002  F (BL vs 300 min): *p* = 0.345 |
| 5E | n = 8 M  n = 9 F | Two-way ANOVA RM with Bonferroni post hoc correction | *p* = 0.999  *F_A_* = *F*_0.05, (1,15)_ = 0.000 | Mauchly's Test of Sphericity  𝜒^2^ = 39.444  𝜈 = 27  *p* = 0.068 (passed)  then,  *p* = 0.865  *F_B_* = *F*_0.05, (7,105)_ = 0.454  *p* = 0.906  *F_AB_* = *F*_0.05, (3.128,50.049)_ = 0.390 | Wilcoxon Signed Rank Test  *𝛂’* = 0.008 | M (BL vs PDSI): *p* = 1.000  M (PDSI vs 30 min): *p* = 0.593  M (30 min vs 60 min): *p* = 0.180  M (60 min vs 120 min): *p* = 0.317  M (120 min vs 180 min): *p* = 0.715  M (180 min vs 240 min): *p* = 0.317  M (240 min vs 300 min): *p* = 0.285  F (BL vs PDSI): *p* = 0.767  F (PDSI vs 30 min): *p* = 0.581  F (30 min vs 60 min): *p* = 0.288  F (60 min vs 120 min): *p* = 1.000  F (120 min vs 180 min): *p* = 0.068  F (180 min vs 240 min): *p* = 0.516  F (240 min vs 300 min): *p* = 0.671 |
|  |  |  |  |  | Wilcoxon Signed Rank Test  *𝛂’* = 0.008 | M (BL vs PDSI):*p* = 1.000  M (BL vs 30 min): *p* = 0.670  M (BL vs 60 min): *p* = 0.888  M (BL vs 120 min): *p* = 0.888  M (BL vs 180 min): *p* = 1.000  M (BL vs 240 min): *p* = 0.673  M (BL vs 300 min): *p* = 1.000  F (BL vs PDSI): *p* = 0.767  F (BL vs 30 min): *p* = 0.859  F (BL vs 60 min): *p* = 0.594  F (BL vs 120 min): *p* = 0.514  F (BL vs 180 min): *p* = 0.514  F (BL vs 240 min): *p* = 0.952  F (BL vs 300 min): *p* = 0.953 |
| 5F | n = 8 M  n = 12 F | Two-way ANOVA RM with Bonferroni post hoc correction | *p* = 0.579  *F_A_* = *F*_0.05, (1,18)_ = 0.319 | Mauchly's Test of Sphericity  𝜒^2^ = 65.367  𝜈 = 27  *p* < 0.001 (failed)  then,  Greenhouse-Geisser:  *p* < 0.001  *F_B_* = *F*_0.05, (4.140,74.524)_ = 56.553 Greenhouse-Geisser:  *p* = 0.806  *F_AB_* = *F*_0.05, (4.140,74.524)_ = 0.412 | Wilcoxon Signed Rank Test  *𝛂’* = 0.008 | M (BL vs PDSI): *p* = 0.011  M (PDSI vs 30 min): *p* = 0.785  M (30 min vs 60 min): *p* = 0.109  M (60 min vs 120 min): *p* = 0.109  M (120 min vs 180 min): *p* = 0.109  M (180 min vs 240 min): *p* = 0.018  M (240 min vs 300 min): *p* = 0.116  F (BL vs PDSI): *p* = 0.002  F (PDSI vs 30 min): *p* = 1.000  F (30 min vs 60 min): *p* = 0.176  F (60 min vs 120 min): *p* = 0.128  F (120 min vs 180 min): *p* = 0.462  F (180 min vs 240 min): *p* = 0.018  F (240 min vs 300 min): *p* = 0.035 |
|  |  |  |  |  | Wilcoxon Signed Rank Test  *𝛂’* = 0.008 | M (BL vs PDSI):*p* = 0.011  M (BL vs 30 min): *p* = 0.012  M (BL vs 60 min): *p* = 0.012  M (BL vs 120 min): *p* = 0.012  M (BL vs 180 min): *p* = 0.012  M (BL vs 240 min): *p* = 0.017  M (BL vs 300 min): *p* = 0.122  F (BL vs PDSI): *p* = 0.002  F (BL vs 30 min): *p* = 0.002  F (BL vs 60 min): *p* = 0.002  F (BL vs 120 min): *p* = 0.002  F (BL vs 180 min): *p* = 0.002  F (BL vs 240 min): *p* = 0.003  F (BL vs 300 min): *p* = 0.181 |

Notes: Two-way ANOVA repeated measure (RM) with Bonferroni post hoc correction was used for this time course data. For within-time two-way ANOVA RM, Greenhouse-Geisser method was used if Mauchly's test of sphericity failed (𝜒^2^, 𝜈, *p*, and *𝛂*’ were values for calculation, degree of freedom (df), significance from Mauchly's test of sphericity and correction for *𝛂* in paired comparisons). *F_A_* was a value for between-subjects ANOVA, *F_B_* was a value for within-subjects ANOVA, while *F_AB_* was a value for interaction effects ANOVA. For paired comparison between each time points, Wilcoxon signed rank test was used if Shapiro-Wilk test and Equal variance test failed. *p* < 0.05 was considered as statistically significant. M, male; F, female.
