## Supplemental Table SIII for "Empathic contagious pain and consolation in laboratory rodents: species and sex comparisons"

Table SIII Sex comparisons of stimulus-response functional curves in mice

| Figure number | Number of animals | Statistic method 1 | Between-sample effects | Within-sample effects | Statistic method 2 | *P* values |
| --- | --- | --- | --- | --- | --- | --- |
| 4A | n = 12 M  n = 12 Mbaseline | Two-way ANOVA RM with Bonferroni post hoc correction | *p* = 0.796  *F_A_* = *F*_0.05, (1,22)_ = 0.068 | Mauchly's Test of Sphericity  𝜒^2^ = 8.137  𝜈 = 9  *p* = 0.522 (passed)  then,  *p* < 0.001  *F_B_* = *F*_0.05, (4,88)_ = 241.349  *p* = 0.628  *F_AB_* = *F*_0.05, (4,88)_ = 0.651 | Friedman′s *M* test | M: *p* < 0.001  Mbaseline: *p* < 0.001 |
|  |  |  |  |  | two-tailed M-W *U* test | M vs Mbaseline (0.16 g): *p* = 0.755  M vs Mbaseline (0.40 g): *p* = 0.755  M vs Mbaseline (0.60 g): *p* = 0.551  M vs Mbaseline (1.00 g): *p* = 0.551  M vs Mbaseline (1.40 g): *p* = 0.630 |
|  | n = 12 F  n =12 Fbaseline | Two-way ANOVA RM with Bonferroni post hoc correction | *p* = 0.509  *F_A_* = *F*_0.05, (1,22)_ = 0.450 | Mauchly's Test of Sphericity  𝜒^2^ = 7.827  𝜈 = 9  *p* = 0.553 (passed)  then,  *p* < 0.001  *F_B_* = *F*_0.05, (4,88)_ = 355.667  *p* = 0.454  *F_AB_* = *F*_0.05, (4,88)_ = 0.923 | Friedman′s *M* test | F: *p* < 0.001  Fbaseline: *p* < 0.001 |
|  |  |  |  |  | two-tailed M-W *U* test | F vs Fbaseline (0.16 g): *p* = 0.319  F vs Fbaseline (0.40 g): *p* = 0.630  F vs Fbaseline (0.60 g): *p* = 0.755  F vs Fbaseline (1.00 g): *p* = 0.551  F vs Fbaseline (1.40 g): *p* = 0.799 |
| 4B | n = 12 M  n = 12 M_baseline_ | Two-way ANOVA RM with Bonferroni post hoc correction | *p* = 0.210  *F_A_* = *F*_0.05, (1,22)_ = 1.669 | Mauchly's Test of Sphericity  𝜒^2^ = 10.173  𝜈 = 9  *p* = 0.338 (passed)  then,  *p* < 0.001  *F_B_* = *F*_0.05, (4,88)_ = 239.682  *p* = 0.832  *F_AB_* = *F*_0.05, (4,88)_ = 0.366 | Friedman′s *M* test | M: *p* < 0.001 |
|  |  |  |  |  | two-tailed M-W *U* test | M vs Mbaseline (0.16 g): *p* = 0.514  M vs Mbaseline (0.40 g): *p* = 0.551  M vs Mbaseline (0.60 g): *p* = 0.551  M vs Mbaseline (1.00 g): *p* = 0.799  M vs Mbaseline (1.40 g): *p* = 0.551 |
|  | n = 12 F  n = 12 Fbaseline | Two-way ANOVA RM with Bonferroni post hoc correction | *p* = 0.885  *F_A_* = *F*_0.05, (1,22)_ = 0.021 | Mauchly's Test of Sphericity  𝜒^2^ = 13.334  𝜈 = 9  *p* = 0.149 (passed)  then,  *p* < 0.001  *F_B_* = *F*_0.05, (4,88)_ = 349.619  *p* = 0.368  *F_AB_* = *F*_0.05, (4,88)_ = 1.087 | Friedman′s *M* test | F: *p* < 0.001 |
|  |  |  |  |  | two-tailed M-W *U* test | F vs Fbaseline (0.16 g): *p* = 0.319  F vs Fbaseline (0.40 g): *p* = 0.590  F vs Fbaseline (0.60 g): *p* = 1.000  F vs Fbaseline (1.00 g): *p* = 1.000  F vs Fbaseline (1.40 g): *p* = 0.590 |
| 4C | n = 12 M  n = 12 Mbaseline | Two-way ANOVA RM with Bonferroni post hoc correction | *p* = 0.328  *F_A_* = *F*_0.05, (1,22)_ = 1.000 | Mauchly's Test of Sphericity  𝜒^2^ = 10.271  𝜈 = 9  *p* = 0.331 (passed)  then,  *p* < 0.001  *F_B_* = *F*_0.05, (4,88)_ = 281.095  *p* = 0.422  *F_AB_* = *F*_0.05, (4,88)_ = 0.981 | Friedman′s *M* test | M: *p* < 0.001 |
|  |  |  |  |  | two-tailed M-W *U* test | M vs Mbaseline (0.16 g): *p* = 0.755  M vs Mbaseline (0.40 g): *p* = 1.000  M vs Mbaseline (0.60 g): *p* = 1.000  M vs Mbaseline (1.00 g): *p* = 0.242  M vs Mbaseline (1.40 g): *p* = 0.378 |
|  | n = 12 F  n = 12 Fbaseline | Two-way ANOVA RM with Bonferroni post hoc correction | *p* = 0.770  *F_A_* = *F*_0.05, (1,22)_ = 0.088 | Mauchly's Test of Sphericity  𝜒^2^ = 11.897  𝜈 = 9  *p* = 0.221 (passed)  then,  *p* < 0.001  *F_B_* = *F*_0.05, (4,88)_ = 327.442  *p* = 0.148  *F_AB_* = *F*_0.05, (4,88)_ = 1.740 | Friedman′s *M* test | F: *p* < 0.001 |
|  |  |  |  |  | two-tailed M-W *U* test | F vs Fbaseline (0.16 g): *p* = 0.514  F vs Fbaseline (0.40 g): *p* = 0.590  F vs Fbaseline (0.60 g): *p* = 0.551  F vs Fbaseline (1.00 g): *p* = 1.000  F vs Fbaseline (1.40 g): *p* = 0.219 |
| 4D | n = 12 M  n = 12 Mbaseline | Two-way ANOVA RM with Bonferroni post hoc correction | *p* = 0.758  *F_A_* = *F*_0.05, (1,22)_ = 0.097 | Mauchly's Test of Sphericity  𝜒^2^ = 7.131  𝜈 = 9  *p* = 0.625 (passed)  then,  *p* < 0.001  *F_B_* = *F*_0.05, (2.555,56.201)_ = 349.852  *p* = 0.987  *F_AB_* = *F*_0.05, (2.555,56.201)_ = 0.086 | Friedman′s *M* test | M: *p* < 0.001 |
|  |  |  |  |  | two-tailed M-W *U* test | M vs Mbaseline (0.16 g): *p* = 1.000  M vs Mbaseline (0.40 g): *p* = 0.799  M vs Mbaseline (0.60 g): *p* = 1.000  M vs Mbaseline (1.00 g): *p* = 0.977  M vs Mbaseline (1.40 g): *p* = 0.799 |
|  | n = 12 F  n = 12 Fbaseline | Two-way ANOVA RM with Bonferroni post hoc correction | *p* = 0.852  *F_A_* = *F*_0.05, (1,22)_ = 0.036 | Mauchly's Test of Sphericity  𝜒^2^ = 20.223  𝜈 = 9  *p* = 0.017 (failed)  then,  *p* < 0.001  *F_B_* = *F*_0.05, (2.611,57.435)_ = 350.454  *p* = 0.705  *F_AB_* = *F*_0.05, (2.611,57.435)_ = 0.431 | Friedman′s *M* test | F: *p* < 0.001 |
|  |  |  |  |  | two-tailed M-W *U* test | F vs Fbaseline (0.16 g): *p* = 0.755  F vs Fbaseline (0.40 g): *p* = 1.000  F vs Fbaseline (0.60 g): *p* = 0.799  F vs Fbaseline (1.00 g): *p* = 0.755  F vs Fbaseline (1.40 g): *p* = 0.514 |
| 4E | n = 12 M  n = 12 Mbaseline | Two-way ANOVA RM with Bonferroni post hoc correction | *p* = 0.383  *F_A_* = *F*_0.05, (1,22)_ = 0.793 | Mauchly's Test of Sphericity  𝜒^2^ = 5.579  𝜈 = 9  *p* = 0.782 (passed)  then,  *p* < 0.001  *F_B_* = *F*_0.05, (4,88)_ = 318.580  *p* = 0.457  *F_AB_* = *F*_0.05, (4,88)_ = 0.919 | Friedman′s *M* test | M: *p* < 0.001 |
|  |  |  |  |  | two-tailed M-W *U* test | M vs Mbaseline (0.16 g): *p* = 0.755  M vs Mbaseline (0.40 g): *p* = 0.551  M vs Mbaseline (0.60 g): *p* = 0.755  M vs Mbaseline (1.00 g): *p* = 0.551  M vs Mbaseline (1.40 g): *p* = 0.347 |
|  | n = 12 F  n = 12 Fbaseline | Two-way ANOVA RM with Bonferroni post hoc correction | *p* = 0.510  *F_A_* = *F*_0.05, (1,22)_ = 0.449 | Mauchly's Test of Sphericity  𝜒^2^ = 9.129  𝜈 = 9  *p* = 0.427 (passed)  then,  *p* < 0.001  *F_B_* = *F*_0.05, (4,88)_ = 357.872  *p* = 0.782  *F_AB_* = *F*_0.05, (4,88)_ = 0.436 | Friedman′s *M* test | F: *p* < 0.001 |
|  |  |  |  |  | two-tailed M-W *U* test | F vs Fbaseline (0.16 g): *p* = 0.755  F vs Fbaseline (0.40 g): *p* = 0.590  F vs Fbaseline (0.60 g): *p* = 0.755  F vs Fbaseline (1.00 g): *p* = 1.000  F vs Fbaseline (1.40 g): *p* = 0.551 |
| 4F | n = 12 M  n = 12 Mbaseline | Two-way ANOVA RM with Bonferroni post hoc correction | *p* = 0.430  *F_A_* = *F*_0.05, (1,22)_ = 0.647 | Mauchly's Test of Sphericity  𝜒^2^ = 7.488  𝜈 = 9  *p* = 0.588 (passed)  then,  *p* < 0.001  *F_B_* = *F*_0.05, (4,88)_ = 342.640  *p* = 0.750  *F_AB_* = *F*_0.05, (4,88)_ = 0.481 | Friedman′s *M* test | M: *p* < 0.001 |
|  |  |  |  |  | two-tailed M-W *U* test | M vs Mbaseline (0.16 g): *p* = 0.755  M vs Mbaseline (0.40 g): *p* = 1.000  M vs Mbaseline (0.60 g): *p* = 0.347  M vs Mbaseline (1.00 g): *p* = 0.551  M vs Mbaseline (1.40 g): *p* = 1.000 |
|  | n = 12 F  n = 12 Fbaseline | Two-way ANOVA RM with Bonferroni post hoc correction | *p* = 0.681  *F_A_* = *F*_0.05, (1,22)_ = 0.173 | Mauchly's Test of Sphericity  𝜒^2^ = 3.261  𝜈 = 9  *p* = 0.953 (passed)  then,  *p* < 0.001  *F_B_* = *F*_0.05, (4,88)_ = 262.809  *p* = 0.079  *F_AB_* = *F*_0.05, (4,88)_ = 2.170 | Friedman′s *M* test | F: *p* < 0.001 |
|  |  |  |  |  | two-tailed M-W *U* test | F vs Fbaseline (0.16 g): *p* = 0.178  F vs Fbaseline (0.40 g): *p* = 0.799  F vs Fbaseline (0.60 g): *p* = 0.590  F vs Fbaseline (1.00 g): *p* = 0.478  F vs Fbaseline (1.40 g): *p* = 0.128 |
| 4G | n = 12 M  n = 12 Mbaseline | Two-way ANOVA RM with Bonferroni post hoc correction | *p* = 0.019  *F_A_* = *F*_0.05, (1,22)_ = 6.456 | Mauchly's Test of Sphericity  𝜒^2^ = 10.301  𝜈 = 9  *p* = 0.328 (passed)  then,  *p* < 0.001  *F_B_* = *F*_0.05, (4,88)_ = 181.462  *p* = 0.231  *F_AB_* = *F*_0.05, (4,88)_ = 1.429 | Friedman′s *M* test | M: *p* < 0.001  Mbaseline: *p* < 0.001 |
|  |  |  |  |  | two-tailed M-W *U* test | M vs Mbaseline (0.16 g): *p* = 0.024  M vs Mbaseline (0.40 g): *p* = 0.378  M vs Mbaseline (0.60 g): *p* = 0.089  M vs Mbaseline (1.00 g): *p* = 0.020  M vs Mbaseline (1.40 g): *p* = 0.114 |
|  | n = 12 F  n =12 Fbaseline | Two-way ANOVA RM with Bonferroni post hoc correction | *p* < 0.001  *F_A_* = *F*_0.05, (1,22)_ = 84.666 | Mauchly's Test of Sphericity  𝜒^2^ = 8.718  𝜈 = 9  *p* = 0.465 (passed)  then,  *p* < 0.001  *F_B_* = *F*_0.05, (4,88)_ = 318.449  *p* = 0.060  *F_AB_* = *F*_0.05, (4,88)_ = 2.349 | Friedman′s *M* test | F: *p* < 0.001  Fbaseline: *p* < 0.001 |
|  |  |  |  |  | two-tailed M-W *U* test | F vs Fbaseline (0.16 g): *p* = 0.178  F vs Fbaseline (0.40 g): *p* < 0.001  F vs Fbaseline (0.60 g): *p* < 0.001  F vs Fbaseline (1.00 g): *p* < 0.001  F vs Fbaseline (1.40 g): *p* = 0.002 |
| 4H | n = 12 M  n = 12 M_baseline_ | Two-way ANOVA RM with Bonferroni post hoc correction | *p* < 0.001  *F_A_* = *F*_0.05, (1,22)_ = 16.656 | Mauchly's Test of Sphericity  𝜒^2^ = 22.759  𝜈 = 9  *p* = 0.007 (failed)  then,  Greenhouse-Geisser:  *p* < 0.001  *F_B_* = *F*_0.05, (2.393,52.643)_ = 129.761  Greenhouse-Geisser:  *p* = 0.088  *F_AB_* = *F*_0.05, (2.393,52.643)_ = 2.434 | Friedman′s *M* test | M: *p* < 0.001 |
|  |  |  |  |  | two-tailed M-W *U* test | M vs Mbaseline (0.16 g): *p* < 0.001  M vs Mbaseline (0.40 g): *p* = 0.007  M vs Mbaseline (0.60 g): *p* = 0.014  M vs Mbaseline (1.00 g): *p* = 0.028  M vs Mbaseline (1.40 g): *p* = 0.012 |
|  | n = 12 F  n = 12 Fbaseline | Two-way ANOVA RM with Bonferroni post hoc correction | *p* < 0.001  *F_A_* = *F*_0.05, (1,22)_ = 57.863 | Mauchly's Test of Sphericity  𝜒^2^ = 16.391  𝜈 = 9  *p* = 0.060 (passed)  then,  *p* < 0.001  *F_B_* = *F*_0.05, (4,88)_ = 367.013  *p* = 0.363  *F_AB_* = *F*_0.05, (4,88)_ = 1.097 | Friedman′s *M* test | F: *p* < 0.001 |
|  |  |  |  |  | two-tailed M-W *U* test | F vs Fbaseline (0.16 g): *p* = 0.014  F vs Fbaseline (0.40 g): *p* = 0.003  F vs Fbaseline (0.60 g): *p* < 0.001  F vs Fbaseline (1.00 g): *p* < 0.001  F vs Fbaseline (1.40 g): *p* = 0.007 |
| 4I | n = 12 M  n = 12 Mbaseline | Two-way ANOVA RM with Bonferroni post hoc correction | *p* = 0.003  *F_A_* = *F*_0.05, (1,22)_ = 11.142 | Mauchly's Test of Sphericity  𝜒^2^ = 23.848  𝜈 = 9  *p* = 0.005 (failed)  then,  Greenhouse-Geisser:  *p* < 0.001  *F_B_* = *F*_0.05, (2.474,54.431)_ = 136.651  Greenhouse-Geisser:  *p* = 0.060  *F_AB_* = *F*_0.05, (2.474,54.431)_ = 2.773 | Friedman′s *M* test | M: *p* < 0.001 |
|  |  |  |  |  | two-tailed M-W *U* test | M vs Mbaseline (0.16 g): *p* = 0.003  M vs Mbaseline (0.40 g): *p* = 0.014  M vs Mbaseline (0.60 g): *p* = 0.008  M vs Mbaseline (1.00 g): *p* = 0.028  M vs Mbaseline (1.40 g): *p* = 0.178 |
|  | n = 12 F  n = 12 Fbaseline | Two-way ANOVA RM with Bonferroni post hoc correction | *p* < 0.001  *F_A_* = *F*_0.05, (1,22)_ = 58.482 | Mauchly's Test of Sphericity  𝜒^2^ = 6.200  𝜈 = 9  *p* = 0.721 (passed)  then,  *p* < 0.001  *F_B_* = *F*_0.05, (4,88)_ = 288.422  *p* = 0.505  *F_AB_* = *F*_0.05, (4,88)_ = 0.837 | Friedman′s *M* test | F: *p* < 0.001 |
|  |  |  |  |  | two-tailed M-W *U* test | F vs Fbaseline (0.16 g): *p* = 0.039  F vs Fbaseline (0.40 g): *p* = 0.002  F vs Fbaseline (0.60 g): *p* = 0.001  F vs Fbaseline (1.00 g): *p* < 0.001  F vs Fbaseline (1.40 g): *p* = 0.007 |
| 4J | n = 12 M  n = 12 Mbaseline | Two-way ANOVA RM with Bonferroni post hoc correction | *p* = 0.002  *F_A_* = *F*_0.05, (1,22)_ = 12.571 | Mauchly's Test of Sphericity  𝜒^2^ = 20.277  𝜈 = 9  *p* = 0.017 (failed)  then,  Greenhouse-Geisser:  *p* < 0.001  *F_B_* = *F*_0.05, (2.555,56.201)_ = 124.570  Greenhouse-Geisser:  *p* = 0.024  *F_AB_* = *F*_0.05, (2.555,56.201)_ = 3.624 | Friedman′s *M* test | M: *p* < 0.001 |
|  |  |  |  |  | two-tailed M-W *U* test | M vs Mbaseline (0.16 g): *p* < 0.001  M vs Mbaseline (0.40 g): *p* = 0.005  M vs Mbaseline (0.60 g): *p* = 0.060  M vs Mbaseline (1.00 g): *p* = 0.028  M vs Mbaseline (1.40 g): *p* = 0.178 |
|  | n = 12 F  n = 12 Fbaseline | Two-way ANOVA RM with Bonferroni post hoc correction | *p* < 0.001  *F_A_* = *F*_0.05, (1,22)_ = 59.703 | Mauchly's Test of Sphericity  𝜒^2^ = 6.284  𝜈 = 9  *p* = 0.712 (passed)  then,  *p* < 0.001  *F_B_* = *F*_0.05, (4,88)_ = 336.267  *p* = 0.650  *F_AB_* = *F*_0.05, (4,88)_ = 0.619 | Friedman′s *M* test | F: *p* < 0.001 |
|  |  |  |  |  | two-tailed M-W *U* test | F vs Fbaseline (0.16 g): *p* = 0.014  F vs Fbaseline (0.40 g): *p* = 0.001  F vs Fbaseline (0.60 g): *p* = 0.001  F vs Fbaseline (1.00 g): *p* < 0.001  F vs Fbaseline (1.40 g): *p* = 0.007 |
| 4K | n = 12 M  n = 12 Mbaseline | Two-way ANOVA RM with Bonferroni post hoc correction | *p* = 0.001  *F_A_* = *F*_0.05, (1,22)_ = 16.477 | Mauchly's Test of Sphericity  𝜒^2^ = 31.196  𝜈 = 9  *p* < 0.001(failed)  then,  Greenhouse-Geisser:  *p* < 0.001  *F_B_* = *F*_0.05, (2.201,48.422)_ = 120.232  Greenhouse-Geisser:  *p* = 0.030  *F_AB_* = *F*_0.05, (2.201,48.422)_ = 3.649 | Friedman′s *M* test | M: *p* < 0.001 |
|  |  |  |  |  | two-tailed M-W *U* test | M vs Mbaseline (0.16 g): *p* = 0.002  M vs Mbaseline (0.40 g): *p* = 0.007  M vs Mbaseline (0.60 g): *p* = 0.002  M vs Mbaseline (1.00 g): *p* = 0.020  M vs Mbaseline (1.40 g): *p* = 0.291 |
|  | n = 12 F  n = 12 Fbaseline | Two-way ANOVA RM with Bonferroni post hoc correction | *p* = 0.003  *F_A_* = *F*_0.05, (1,22)_ = 11.355 | Mauchly's Test of Sphericity  𝜒^2^ = 6.286  𝜈 = 9  *p* = 0.712 (passed)  then,  *p* < 0.001  *F_B_* = *F*_0.05, (4,88)_ = 231.595  *p* = 0.697  *F_AB_* = *F*_0.05, (4,88)_ = 0.554 | Friedman′s *M* test | F: *p* < 0.001 |
|  |  |  |  |  | two-tailed M-W *U* test | F vs Fbaseline (0.16 g): *p* = 0.160  F vs Fbaseline (0.40 g): *p* = 0.068  F vs Fbaseline (0.60 g): *p* = 0.089  F vs Fbaseline (1.00 g): *p* = 0.068  F vs Fbaseline (1.40 g): *p* = 0.551 |
| 4L | n = 12 M  n = 12 Mbaseline | Two-way ANOVA RM with Bonferroni post hoc correction | *p* = 0.014  *F_A_* = *F*_0.05, (1,22)_ = 7.061 | Mauchly's Test of Sphericity  𝜒^2^ = 21.772  𝜈 = 9  *p* = 0.010 (failed)  then,  Greenhouse-Geisser:  *p* < 0.001  *F_B_* = *F*_0.05, (2.571,56.557)_ = 129.489  Greenhouse-Geisser:  *p* = 0.123  *F_AB_* = *F*_0.05, (2.571,56.557)_ = 2.070 | Friedman′s *M* test | M: *p* < 0.001 |
|  |  |  |  |  | two-tailed M-W *U* test | M vs Mbaseline (0.16 g): *p* = 0.002  M vs Mbaseline (0.40 g): *p* = 0.014  M vs Mbaseline (0.60 g): *p* = 0.319  M vs Mbaseline (1.00 g): *p* = 0.178  M vs Mbaseline (1.40 g): *p* = 0.291 |
|  | n = 12 F  n = 12 Fbaseline | Two-way ANOVA RM with Bonferroni post hoc correction | *p* = 0.001  *F_A_* = *F*_0.05, (1,22)_ = 13.497 | Mauchly's Test of Sphericity  𝜒^2^ = 12.877  𝜈 = 9  *p* = 0.170 (passed)  then,  *p* < 0.001  *F_B_* = *F*_0.05, (4,88)_ = 261.695  *p* = 0.685  *F_AB_* = *F*_0.05, (4,88)_ = 0.571 | Friedman′s *M* test | F: *p* < 0.001 |
|  |  |  |  |  | two-tailed M-W *U* test | F vs Fbaseline (0.16 g): *p* = 0.089  F vs Fbaseline (0.40 g): *p* = 0.178  F vs Fbaseline (0.60 g): *p* = 0.319  F vs Fbaseline (1.00 g): *p* = 0.068  F vs Fbaseline (1.40 g): *p* = 0.551 |

Notes: Two-way ANOVA repeated measure (RM) with Bonferroni post hoc correction was used for this time course data. For within-time two-way ANOVA RM, Greenhouse-Geisser method was used if Mauchly's test of sphericity failed (𝜒^2^, 𝜈, and *p* were values for calculation, degree of freedom (df), and significance from Mauchly's test of sphericity). *F_A_* was a value for between-subjects ANOVA, *F_B_* was a value for within-subjects ANOVA, while *F_AB_* was a value for interaction effects ANOVA. For paired comparison between each vF intensity, Friedman′s *M* test and Mann-Whitney *U* test (two-tailed) were used if Shapiro-Wilk test and Equal variance test failed. *p* < 0.05 was considered as statistically significant. M, male; Mbaseline, malebaseline; F, female; Fbaseline, femalebaseline; M-W *U* test, Mann-Whitney *U* test.
