## Supplemental Table SIV for "Empathic contagious pain and consolation in laboratory rodents: species and sex comparisons"

**Table SIV. Sample size prediction by one-way ANOVA Power Analysis**

| **Group of study** | **Sample size predicted** | **95% CI** | **Sample size used** | ***P* value** |
| --- | --- | --- | --- | --- |
| Consolation in mice | Fig 3A  Power = 0.97626, n = 2.00, k = 2, N = 4, Alpha = 0.0500, Beta = 0.02374, Sm = 25.75, SD = 5.92, Effect Size = 4.3497 | (2.33,8.67) | 12 | two-tailed M-W *U* test  *p* = 0.002 |
|  | Fig 3B  Power = 0.92087, n = 9.00, k = 2, N = 18, Alpha = 0.0500, Beta = 0.07913, Sm = 3.30, SD = 3.89, Effect Size = 0.8470 |  |  |  |
|  | Fig 3C  Power = 0.92469, n = 4.00, k = 2, N = 8, Alpha = 0.0500, Beta = 0.07531, Sm = 2.88, SD = 1.98, Effect Size = 1.4520 |  |  |  |
|  | Fig 3D  Power = 0.92565, n = 6.00, k = 2, N = 12, Alpha = 0.0500, Beta = 0.07435, Sm = 1.59, SD = 1.45, Effect Size = 1.0931 |  |  |  |
|  | Fig 3E  Power = 0.93082, n = 3.00, k = 2, N = 6, Alpha = 0.0500, Beta = 0.06918, Sm = 6.13, SD = 3.20, Effect Size = 1.9141 |  |  |  |
|  | Fig 3F  Power = 0.90291, n = 9.00, k = 2, N = 18, Alpha = 0.0500, Beta = 0.09709, Sm = 1.67, SD = 2.04, Effect Size = 0.8186 |  |  |  |
| Observational contagious pain in mice | Fig 4K in 0.60 g  Power = 0.90698, n = 7.00, k = 2, N = 14, Alpha = 0.0500, Beta = 0.09302, Sm = 10.00, SD = 10.45, Effect Size = 0.9569 | (2.60,14.07) | 12 | two-tailed M-W *U* test  *p* = 0.100 |
|  | Fig 4K in 1.00 g  Power = 0.92440, n = 11.00, k = 2, N = 22, Alpha = 0.0500, Beta = 0.07560, Sm = 7.50, SD = 9.85, Effect Size = 0.7614 |  |  |  |
|  | Fig 5B in 240 min  Power = 0.92907, n = 7.00, k = 2, N = 14, Alpha = 0.0500, Beta = 0.07093, Sm = 0.13, SD = 0.13, Effect Size = 1.0000 |  |  |  |
| CO_naive_-CD_naive_ vs CO_naive_-CD_pain_ in mice for Table I | Latency of allolicking/-grooming in male  Power = 1.00000, n = 2.00, k = 2, N = 4, Alpha = 0.0500, Beta = 0.00000, Sm = 405.93, SD = 17.57, Effect Size = 23.1033 | (2.48,8.63) | 12 | two-tailed M-W *U* test  *p <* 0.001 |
|  | Total time spent on allolicking/-grooming in male  Power = 1.00000, n = 2.00, k = 2, N = 4, Alpha = 0.0500, Beta = 0.00000, Sm = 36.54, SD = 2.75, Effect Size = 13.3072 |  |  |  |
|  | Counts of allolicking/-grooming in male  Power = 0.99924, n = 2.00, k = 2, N = 4, Alpha = 0.0500, Beta = 0.00076, Sm = 8.29, SD = 1.37, Effect Size = 6.0492 |  |  |  |
|  | Total time spent on allo-mouth sniffing in male  Power = 1.00000, n = 2.00, k = 2, N = 4, Alpha = 0.0500, Beta = 0.00000, Sm = 445.50, SD = 27.28, Effect Size = 16.3306 |  |  |  |
|  | Total time spent on allo-mouth sniffing in male  Power = 0.90334, n = 12.00, k = 2, N = 24, Alpha = 0.0500, Beta = 0.09666, Sm = 2.46, SD = 3.53, Effect Size = 0.6967 |  |  |  |
|  | Counts of allo-mouth sniffing in male  Power = 0.93902, n = 8.00, k = 2, N = 16, Alpha = 0.0500, Beta = 0.06098, Sm = 1.46, SD = 1.55, Effect Size = 0.9440 |  |  |  |
|  | Total time spent on allo-tail sniffing in male  Power = 0.91495, n = 11.00, k = 2, N = 22, Alpha = 0.0500, Beta = 0.08505, Sm = 5.00, SD = 6.69, Effect Size = 0.7472 |  |  |  |
|  | Total time spent on self-licking/-grooming in male  Power = 0.93566, n = 6.00, k = 2, N = 12, Alpha = 0.0500, Beta = 0.93566, Sm = 42.31, SD = 37.86, Effect Size = 1.1175 |  |  |  |
|  | Counts of self-licking/-grooming in male  Power = 0.95249, n = 5.00, k = 2, N = 10, Alpha = 0.0500, Beta = 0.04751, Sm = 3.04, SD = 2.31, Effect Size = 1.3172 |  |  |  |
|  | Latency of allolicking/-grooming in female  Power = 1.00000, n = 2.00, k = 2, N = 4, Alpha = 0.0500, Beta = 0.00000, Sm = 392.24, SD = 13.31, Effect Size = 29.4696 | (1.29,3.51) | 12 | two-tailed M-W *U* test  *p =* 0.008 |
|  | Total time spent on allolicking/-grooming in female  Power = 0.99805, n = 2.00, k = 2, N = 4, Alpha = 0.0500, Beta = 0.00195, Sm = 10.83, SD = 1.92, Effect Size = 5.6336 |  |  |  |
|  | Counts of allolicking/-grooming in female  Power = 0.97215, n = 2.00, k = 2, N = 4, Alpha = 0.0500, Beta = 0.02785, Sm = 4.96, SD = 1.17, Effect Size = 4.2545 |  |  |  |
|  | Latency of allo-mouth sniffing in female  Power = 1.00000, n = 2.00, k = 2, N = 4, Alpha = 0.0500, Beta = 0.00000, Sm = 506.59, SD = 44.45, Effect Size = 11.3967 |  |  |  |
|  | Latency of self-licking/-grooming in female  Power = 0.96630, n = 4.00, k = 2, N = 4, Alpha = 0.0500, Beta = 0.03370, Sm = 137.75, SD = 84.74, Effect Size = 1.6256 |  |  |  |
| COnaive-CDnaive vs COnaive-CDpain in rats for Table II | Latency of allolicking/-grooming in male  Power = 0.92723, n = 6.00, k = 2, N = 12, Alpha = 0.0500, Beta = 0.07277, Sm = 105.40, SD = 96.10, Effect Size = 1.0968 | (-1.50,8.84) | 8 for CO_naive_-CD_naive_  12 for CO_naive_-CD_pain_ | two-tailed M-W *U* test  CO_naive_-CD_naive_: *p =* 0.100  CO_naive_-CD_pain_: *p =* 0.100 |
|  | Total time spent on allolicking/-grooming in male  Power = 0.99981, n = 2.00, k = 2, N = 4, Alpha = 0.0500, Beta = 0.00019, Sm = 27.90, SD = 4.22, Effect Size = 6.6045 |  |  |  |
|  | Counts of allolicking/-grooming in male  Power = 0.98265, n = 3.00, k = 2, N = 6, Alpha = 0.0500, Beta = 0.01735, Sm = 4.90, SD = 2.13, Effect Size = 2.2950 |  |  |  |
|  | Total time spent on allolicking/-grooming in female  Power = 0.99959, n = 2.00, k = 2, N = 4, Alpha = 0.0500, Beta = 0.00041, Sm = 20.84, SD = 3.31, Effect Size = 6.3011 | (0.90,3.77) | 9 for CO_naive_-CD_naive_  11 for CO_naive_-CD_pain_ | two-tailed M-W *U* test  CO_naive_-CD_naive_: *p =* 0.100  CO_naive_-CD_pain_: *p =* 0.100 |
|  | Counts of allolicking/-grooming in female  Power = 0.99773, n = 3.00, k = 2, N = 6, Alpha = 0.0500, Beta = 0.00227, Sm = 3.88, SD = 1.41, Effect Size = 2.7429 |  |  |  |
|  | Latency of self-licking/-grooming in female  Power = 0.99337, n = 2.00, k = 2, N = 4, Alpha = 0.0500, Beta = 0.00663, Sm = 125.09, SD = 24.79, Effect Size = 5.0460 |  |  |  |

**Notes:** In a one-way ANOVA study, sample sizes of predictions are obtained from the 2 groups whose means are to be compared. The total sample of N subjects achieves power (1 - Beta) to detect differences among the means versus the alternative of equal means using an *F* test with a 0.05 significance level. The size of the variation in the means is represented by their standard deviation: Sm. The common standard deviation within a group is assumed to be SD.

**Report Definitions**

Power is the probability of rejecting a false null hypothesis. It should be close to one.

n is the average group sample size.

k is the number of groups.

Total N is the total sample size of all groups combined.

Alpha is the probability of rejecting a true null hypothesis. It should be small.

Beta is the probability of accepting a false null hypothesis. It should be small.

Sm is the standard deviation of the group means under the alternative hypothesis.

Standard deviation (SD) is the within group standard deviation.

The Effect Size is the ratio of Sm to standard deviation.

M-W *U* test is the Mann-Whitney *U* test.

Fleiss, Joseph L. 1986. The Design and Analysis of Clinical Experiments. John Wiley & Sons. New York.

Kirk, Roger E. 1982. Experimental Design: Procedures for the Behavioral Sciences. Brooks/Cole. Pacific Grove, California.
